# Sialoglycan binding triggers spike opening in a human coronavirus

**DOI:** 10.1101/2023.04.20.536837

**Authors:** Matti F. Pronker, Robert Creutznacher, Ieva Drulyte, Ruben J.G. Hulswit, Zeshi Li, Frank J.M. van Kuppeveld, Joost Snijder, Yifei Lang, Berend-Jan Bosch, Geert-Jan Boons, Martin Frank, Raoul J. de Groot, Daniel L. Hurdiss

## Abstract

Coronavirus (CoV) spikes mediate receptor binding and membrane fusion, making them prime targets for neutralising antibodies. In the cases of SARS-CoV, SARS-CoV-2, and MERS-CoV, spikes transition freely between open and closed conformations to balance host cell attachment and immune evasion. The open conformation exposes domain S1^B^, allowing it to bind to proteinaceous cell surface receptors. It also facilitates protein refolding during spike-mediated membrane fusion. However, with a single exception, the pre-fusion spikes of all other CoVs studied so far have been observed exclusively in the closed state. This raises the possibility of regulation, where spikes more commonly transition to open states in response to specific cues, rather than spontaneously. In our study, using cryo-EM and molecular dynamics simulations, we show that the spike protein of the common cold human coronavirus HKU1 undergoes local and long-range conformational changes upon binding a sialoglycan-based primary receptor to domain S1^A^. This binding triggers the transition of S1^B^ domains to the open state via allosteric inter-domain cross-talk. Our findings paint a more elaborate picture of CoV attachment, with possibilities of dual receptor usage and priming of entry as a means of immune escape.

## INTRODUCTION

Long before the advent of severe acute respiratory syndrome coronavirus 2 (SARS-CoV-2), four coronaviruses (CoVs) colonised the human population. Two of these, human coronaviruses HKU1 and OC43 in the betacoronavirus subgenus *Embecovirus*, independently arose from rodent reservoirs—either directly or via intermediate hosts^1–3^. Unlike other human CoVs, HKU1 and OC43 rely on cell surface glycans as indispensable primary receptors^4, 5^. Their attachment and fusion spike proteins (S) specifically bind to 9-*O*-acetylated sialosides^4, 6–10^. Underlining the importance of glycan attachment, embecoviruses uniquely code for an additional envelope protein, hemagglutinin-esterase, a sialate-*O*-acetylesterase serving as a receptor-destroying enzyme^6, 11, 12^. Recent observations suggest that HKU1 S particularly targets α2,8-linked 9-*O*-acetylated disialosides (9-*O*-Ac-Sia(α2,8)Sia, *i.e.* glycan motifs typical for oligosialogangliosides like GD3. Accordingly, upon overexpression of GD3 synthase ST8SIA1, HEK293T cells become susceptible to HKU1 spike-pseudotyped viruses^10^.

CoV spikes are homo-trimeric class I fusion proteins^13^. The S-protomer can be divided into an N-and C-terminal region designated S1 and S2, respectively. Distinct S1 domains mediate receptor binding^14^, whereas S2 comprises the fusion machinery (Fig. 1a). In HKU1 and OC43, attachment to 9-*O-*Ac-sialosides occurs through a well-conserved receptor binding site located in S protein domain S1^A^ ^8, 9^ (Fig. 1a). There are indications, however, for the existence of a secondary receptor engaged through domain S1^B^, as epitopes of virus-neutralising antibodies map to subdomain S1^B2^ ^15–17^. Moreover, in the case of HKU1, recombinantly expressed S1^B^ blocks infection^16^, with single site substitutions in S1^B^^2^ resulting in loss of this activity^17^.

**Fig. 1.**
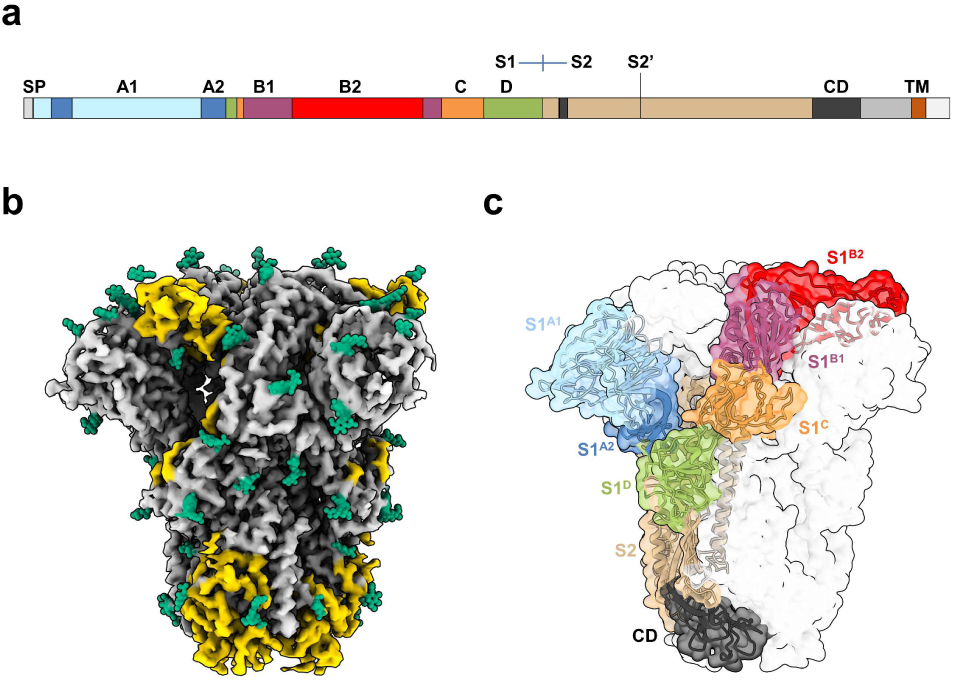
Cryo-EM structure of the *apo* HKU1-A S. **a**, Linear representation of the HKU1-A S primary sequence, coloured by domain. The S1-S2 domains, S2’ protease cleavage site, signal peptide for secretion (SP), connecting domain (CD) and transmembrane helix (TM) are also indicated. **b**, Cryo-EM density map for *apo* HKU1-A S, with previously unmodelled glycans indicated in green. Newly modelled amino acids are coloured yellow. **c**, *Apo* HKU1-A S trimer with one Y-shaped protomer coloured by (sub)domain. The colours correspond to the linear representation of the S domain organisation in **a**.

The S proteins of SARS-CoV, SARS-CoV-2 and MERS-CoV occur in different conformations with their receptor-binding S1^B^ domains either partially buried between neighbouring protomers (‘closed’ or ‘down’) or with one or more S1^B^ domains exposed (1-, 2-and 3-up, ‘open’)^18–21^. The conformational dynamics of S1^B^, and modulation thereof, would provide CoVs with a means to balance host cell attachment and immune escape^22^. Recently, spontaneous conversion of S1^B^ into the up conformation was also described for porcine epidemic diarrhoea virus^23^. Puzzlingly, however, available structures of all other CoV S proteins, including those of HKU1 and OC43^9, 24^, have only been observed in a closed conformation (Extended Data Table 1), shielding S1^B^ from neutralising antibodies but preventing S1^B^-mediated receptor engagement^15, 22^. Adding to the conundrum, the transition from a closed to an open S conformation has been linked to the elaborate conformational changes in S2 that drive fusion^25–27^. The question thus arises whether specific mechanisms might exist that trigger S1^B^ conversion to the open state. Here we describe cryo-EM structures of a serotype A HKU1 (HKU1-A) S in four conformations, one in a closed *apo* state, the others in complex with the HKU1 disialoside receptor 9-*O*-Ac-Sia(α2,8)Sia. We show that glycan receptor binding by S1^A^ specifically prompts a conformational transition of S1^B^ domains into 1-and eventually 3-up positions, apparently through an allosteric mechanism.

## RESULTS

### Cryo-EM structure of the HKU1 type A S glycoprotein

HKU1 field strains are divided into three genotypes with evidence of intertypic recombination, but essentially occur in two distinct serotypes, sporting either A-or B-type spike (S) proteins^28^. Single-particle cryo-EM analysis of S ectodomains of type A HKU1 (HKU1-A) strain Caen1 yielded a reconstruction for the unbound state at a global resolution of 3.4 Å (Fig. 1b, Extended Data Fig. 1-2, Supplementary Table 2). Notably, the HKU1-A S trimers were found exclusively in a closed, pre-fusion conformation as reported for a serotype B HKU1 (HKU1-B) S protein^24^. The HKU1-A and HKU1-B S proteins, at 84% sequence identity (Extended Data Fig. 3), are highly similar in global structure with an average C_α_ RMSD of 1.1 Å for pruned atom pairs (Extended Data Fig. 4). Compared to the HKU1-B model, our data allowed building an additional 231 residues per protomer. Among these newly built segments are the membrane-proximal connecting domain (CD, residues 776-796 and 1152-1225) and the linker between the S1/S2 and S2’ protease cleavage sites (residues 878-907) (Fig. 1c). We could also model a major portion of S1^B^^2^ (residues 480-575) such that this subdomain—purportedly crucial for protein receptor binding—is now fully resolved in the context of an intact HKU1 S trimer, our findings essentially confirming the crystal structure of a HKU1-A S1^B-C^ fragment (residues 310-677)^17^ (Extended Data Fig. 4). In addition, 20 N-linked glycans per protomer were built into the model, all well supported by the density map (Fig. 1b). Several glycans are engaged in interprotomer contacts (*e.g.*, N1215, Extended Data Fig. 5), among which the N355-glycan in S1^B^ may help to stabilise the HKU1-A S trimer in the closed conformation by contacting the clockwise neighbouring protomer via Y528 (Extended Data Fig. 6). Using site-specific glycosylation patterns of HKU1-B^29^, we performed molecular dynamics (MD) simulations of the fully glycosylated S ectodomain trimer, providing a comprehensive overview of exposed and glycan-shielded regions of HKU1-A S (Extended Data Fig. 7).

**Fig. 2.**
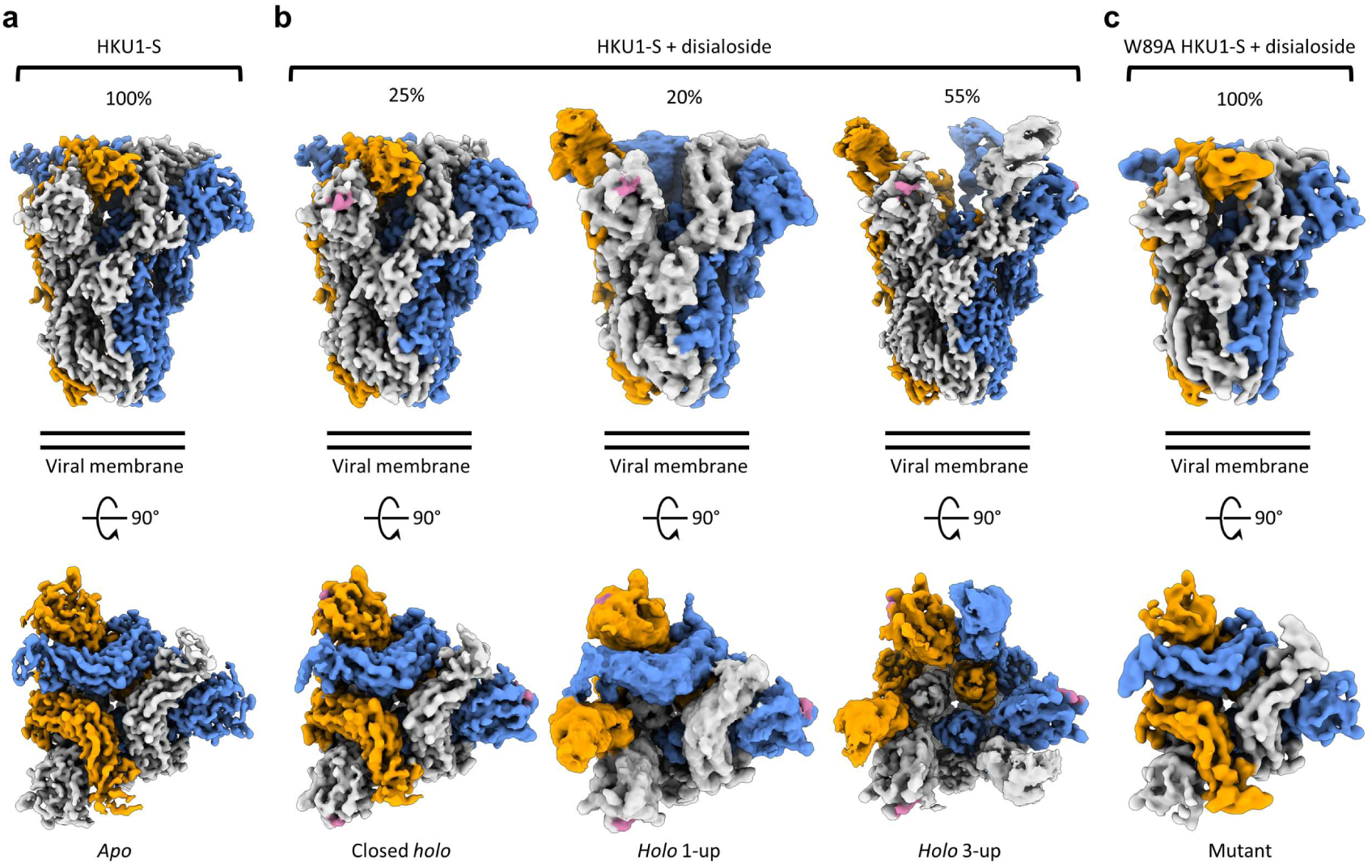
Cryo-EM density maps of wt *apo* HKU1-A S, its complex with a 9-*O*-acetylated disialoside and an equivalently liganded W89A mutant. **a**, Two orthogonal views of the *apo* HKU1-A S trimer density map with protomers coloured in grey, orange, and blue. **b**, Density maps of HKU1-A S in complex with the disialoside. Three distinct classes were observed, with either no, one or three S1^B^ domains in the open conformation. Percentages of particles observed for each reconstruction are indicated above. The bound disialoside is coloured pink. **c**, Cryo-EM density map of the non-binding HKU1-A S W89A mutant in presence of the disialoside has all S1^B^ domains in the closed orientation.

**Fig. 3.**
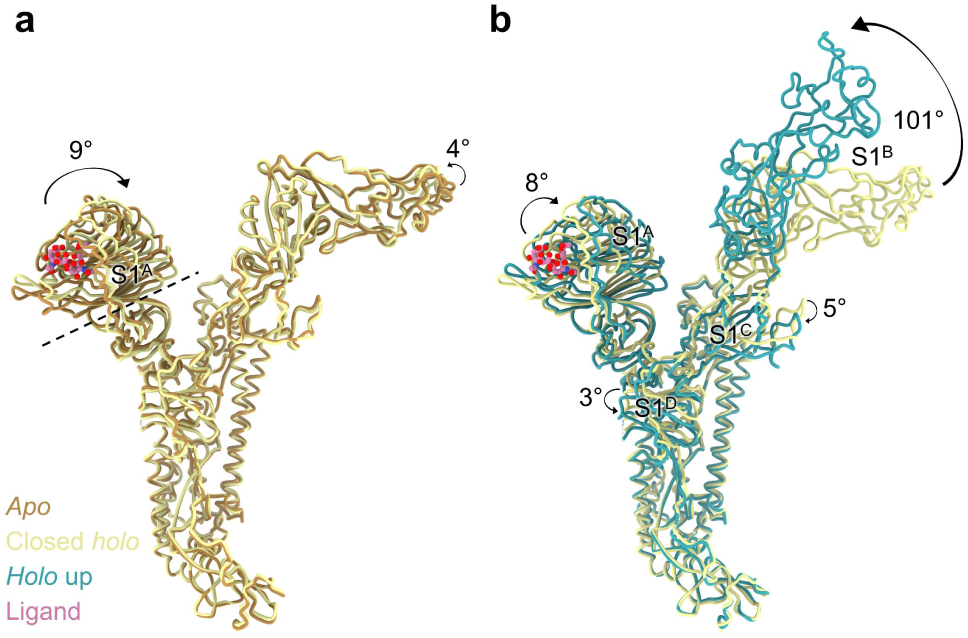
Allosteric inter-domain and intra-domain rotations are observed upon ligand binding. **a**, Superposition of a single protomer of *apo* HKU1-A S with the ligand bound closed *holo* state. The axis around which the S1^A1^ subdomain rotates upon ligand binding is indicated as a dashed line, the disialoside is shown as pink spheres. **b**, Further domain rotations are observed when going from the closed *holo* state to the *holo* up conformation (here from the 3-up state), not only of S1^B^ but also of all other S1 domains.

**Fig. 4.**
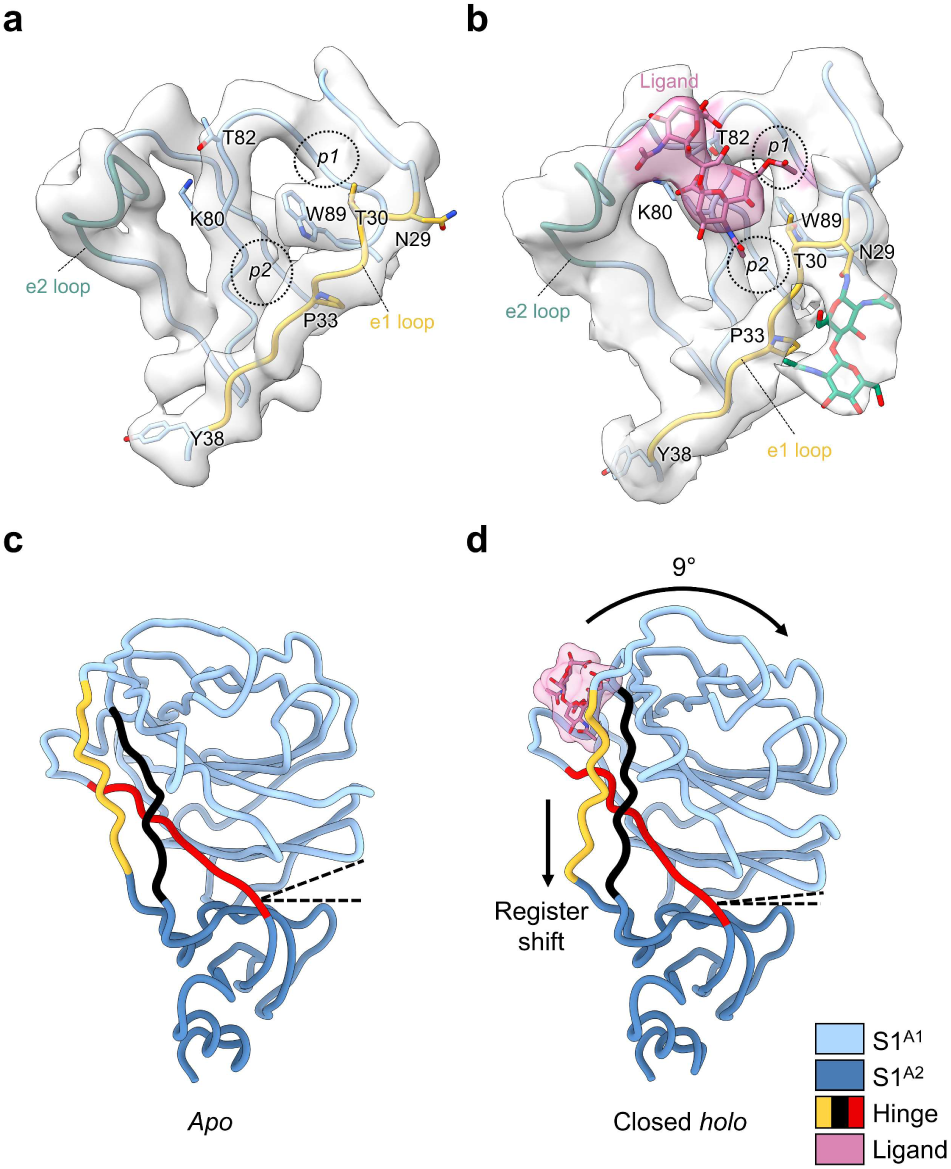
Comparison of the sialic acid binding site in the *apo* and closed *holo* S1^A^ domains. **a**, Sialic acid-binding pocket in the S1^A^ domain in the *apo* state, with the e1 loop indicated in yellow and the e2 loop in green. The *p1* and *p2* pockets are indicated with dashed circles. **b**, The equivalent pocket in the closed *holo* state reveals clear density for the disialoside (pink). The N-linked glycan on N29 becomes ordered, allowing the modelling of two GlcNAc residues (green). Substantial conformational changes are seen in the e1 loop compared to the *apo* state. **c**, S1^A^ domain of one protomer in the *apo* state, coloured by the S1^A1^ (light blue) and S1^A2^ (dark blue) subdomains. The three hinge segments connecting the subdomains are coloured yellow (e1 loop), red and black. The angle between the subdomains is indicated for comparison with the *holo* state in **d**. **d**, Same view of the S1^A^ domain as in **c**, but for the closed *holo* state (disialoside in pink). The angle between the S1^A1^ and S1^A2^ subdomains is notably smaller due to the intradomain inward wedging rotation of the S1^A1^ subdomain.

**Fig. 5.**
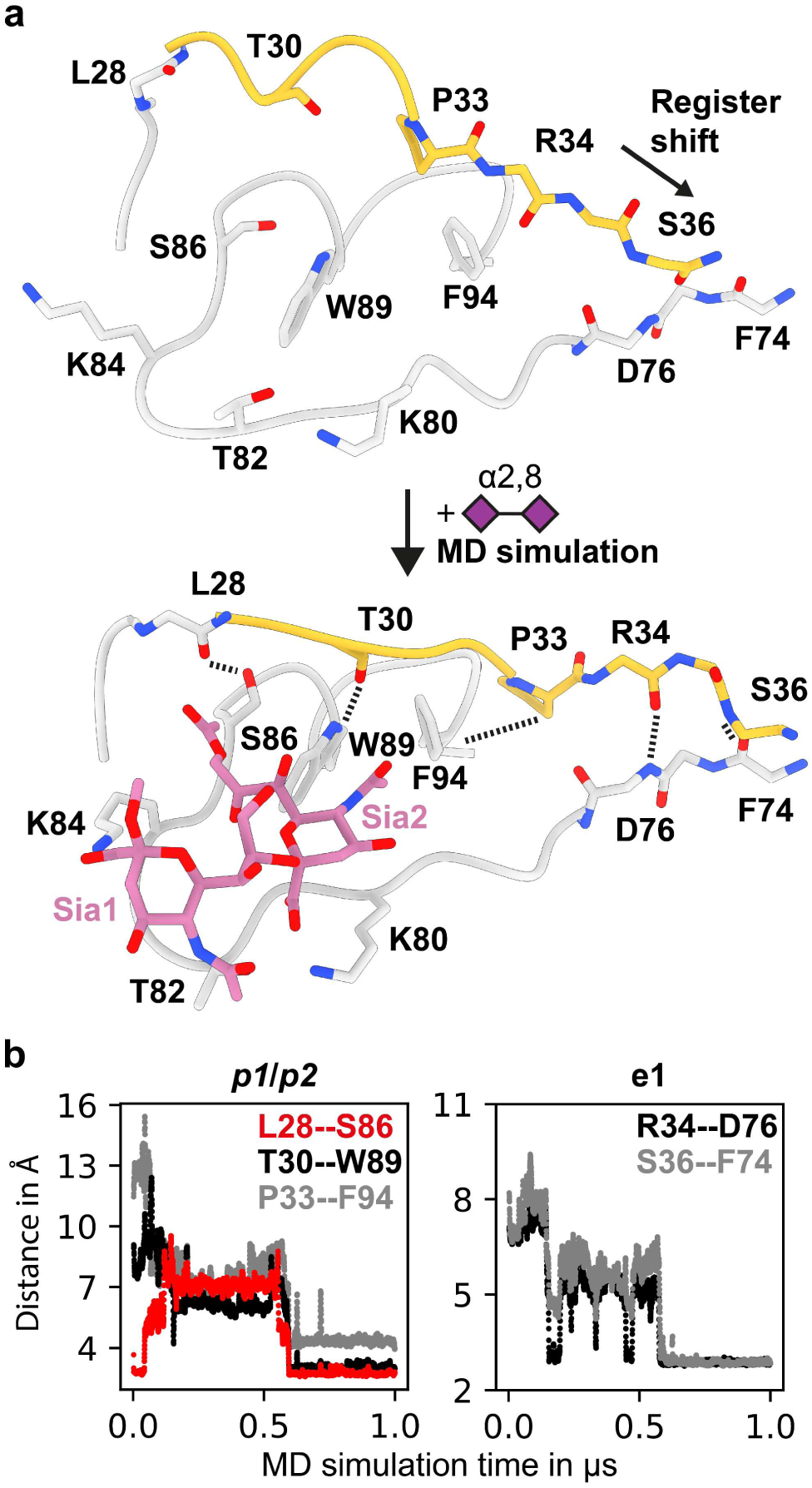
MD simulations predict S1^A^ conformational transition. **a**, An MD trajectory exemplifying that docking the disialoside (symbolic depiction as purple diamonds) into the *apo* cryo-EM model (top), leads to the conformational change into the stable *holo* state (bottom). **b,** Within the e1 loop (yellow), including the *p1* and *p2* pockets, several key hydrogen bonds and hydrophobic contacts form within 500 ns. Changes in inter-residue distances served as a measure to monitor the conformational changes, as the protein transitions into the cryo-EM *holo* state conformation. Only relevant side chains are shown for clarity. The conformational change in **b** is visualised in Supplementary Movie 7.

**Fig. 6.**
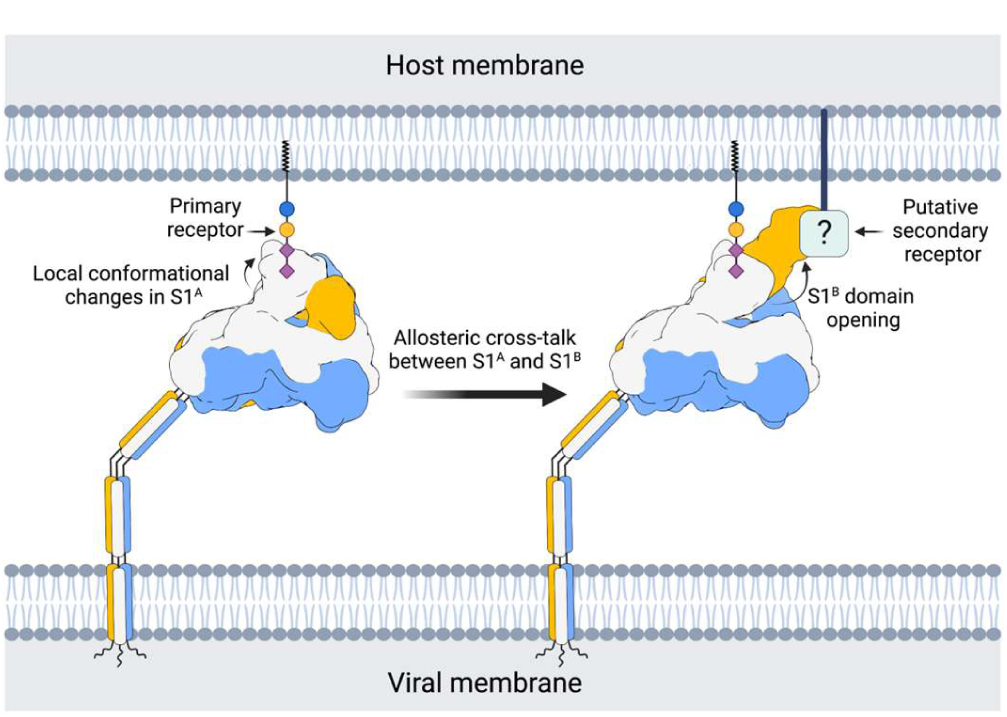
Proposed model for HKU1-A S host cell engagement. Proposed model for HKU1-A S host cell engagement through a primary carbohydrate receptor, containing a 9-*O*-acetylated α2,8-linked disialoside binding to the S1^A^ domain, leading to the allosteric opening of the neighbouring S1^B^ domain and a putative secondary receptor binding to the exposed open S1^B^ domain. Carbohydrates are coloured according to the symbolic nomenclature for glycans^30^.

Predictably similar in their overall arrangement, the *apo* structures of A and B-type S trimers differ in the orientation of their glycan-binding S1^A^ domains, with those of HKU1-A tilted outwards. The S1^A^ 9-*O*-Ac-Sia binding site is conserved in HKU1-A S1^A^, as expected, with key ligand contact residues K80, T/S82, and W89^8^ aligning with those in HKU1-B S (Extended Data Fig. 4, 8). There are, however, notable differences in binding site topology. In HKU1-B, the 9-*O-*Ac-Sia-binding site is located within a narrow crevice between loop elements e1 (residues 29-37) and e2 (residues 246-252)^8, 9^. In the HKU1-A S *apo* structure, the *p1* and *p2* pockets that accommodate the sialoside 9-*O*-and 5-*N*-Ac moieties, respectively, are much less prominent due to a consequential outward displacement of the e1 loop (*vide infra*) (Extended Data Fig. 8).

### Sialoglycan binding triggers allosteric opening of the S1^B^ domain

Incubation of the HKU1-A S protein with the receptor analogue 9-*O*-Ac-Neu5Ac-α2,8-Neu5Ac-Lc-biotin (Extended Data Fig. 9) led to dramatic conformational changes yielding a surprising heterogeneity in structures. We identified and modelled three distinct conformations: a fully closed state (3.8 Å resolution), a partially opened state with a single S1^B^ domain rotated upwards by 101° (1-up, 5 Å resolution), and a fully opened state (3-up, 3.7 Å resolution) (Fig. 2, Extended Data Fig. 10-12, Supplementary Table 2). A 2-up state was not detected. In all *holo* structures, clear densities for the disialoside were observed within S1^A^ receptor binding sites (Fig. 2 and Extended Data Fig. 13-14). Apparently, binding of a specific 9-*O*-Ac-Sia-based primary receptor analogue by the S1^A^ domain triggers an allosteric mechanism, that causes the exposure of S1^B^ domains located 40 Å from the S1^A^ binding pocket (Fig. 2).

Based on our observations, we propose a stepwise model for ligand-induced spike opening (Supplementary Video 1). In the starting *apo* state, each S1^B^ domain is held in place, wedged between the S1^A^ and S1^B^ domains of the counter-clockwise neighbouring Y-shaped protomer. Of the two observed protein-protein interfaces, the one with S1^A^ buries a larger surface area (Extended Data Fig. 6; 1207 Å^2^ versus 442 Å^2^). In the presence of the S1^A^ ligand, most S trimers transitioned into the 1-or 3-up open states. However, 25% of ligand-bound particles remained fully closed. The structure of this ‘closed *holo*’ trimer is distinct from that of non-complexed *apo* trimers, marking it as an initial step in a series of conformational transitions. Ligand binding in the ‘closed *holo*’ state is associated with intradomain conformational changes within S1^A^. In particular, the upper S1^A1^ subdomain (residues 14-39 and 72-260) rotates inwards by 9° relative to S1^A2^ and the remainder of the S monomer (Fig. 3). While this motion leaves the S1^B^-S1^B^ interface unaltered, it has a profound impact on the S1^A^-S1^B^ contact area, displacing interfacing residues by approximately 8 Å (Extended Data Fig. 15). This reshaping of the S1^A^-S1^B^ interface appears to be the key phenomenon from which subsequent upward rotation of the first S1^B^ domain follows, involving a 101° rotation and raising the tip of the S1^B2^ subdomain by 50 Å (Fig. 3b, Supplementary Video 2).

This large conformational change of S1^B^ going from the closed *holo* to the *holo* up state is accompanied by additional domain rotations of S1^A^, S1^C^ and S1^D^ (Fig. 3b). Conversion into the ‘1-up’ state eliminates the S1^B^-S1^B^ interdomain contact (Extended Data Fig. 17). The lack of particles observed in a ‘2-up’ conformation might be explained by the fact that a lone downward oriented S1^B^ lacks any such stabilising interactions with neighbouring S1^B^ domains, likely making this a transient intermediate.

To rule out the possibility that a subset of open S1^B^ domains exist within the *apo* data set, we symmetry-expanded the particles from the *apo* reconstruction and performed 3D variability analysis on the masked S1^B^ domain. No open S1^B^ domains in the *apo* dataset were identified. When the same analysis was performed on the ‘1-up’ particles, open and closed domains could be easily discriminated, confirming the validity of this approach (Extended Data Fig. 18). To further substantiate our observations, we acquired a data set with a sialoglycan binding-defective mutant HKU1 S W89A^8^ in the presence of the 9-*O*-Ac-disialoside as a negative control (Fig. 2c, Extended Data Fig.19-20, Extended Data Table 2). Again, the S trimers were all fully closed and were morphologically indistinguishable from the unbound *apo* state of parental S, reinforcing the notion that binding of 9-*O*-Ac-Sia(α2,8)Sia is key for allosteric release of S1^B^.

### Sialoglycan binding stabilises an alternative conformation in an S1^A^ loop

Local refinement of the symmetrical closed structure of the HKU1-ligand complex allowed us to visualise the disialoside bound in the S1^A^ receptor binding site (Fig. 4, Extended Data Fig. 13, Supplementary Table 3), with both Sia moieties well discernible. The location of the essential terminal Sia (Sia2) is as expected for a canonical 9-*O*-Ac-Sia binding site (Hulswit et al., 2019) and matches that of the *holo* cryo-EM structure of OC43 S^9^ (Extended Data Fig. 4). Its assigned orientation positions the sialate-9-*O*-acetyl and -5-*N*-acetyl moieties so that they can dock into pockets *p1* and *p2*, respectively, astride the perpendicularly placed W89 sidechain. The Sia2 carboxylate is poised to interact with K80 and T82 through a salt bridge and hydrogen bond (Fig. 4b). Using dedicated MD simulations of the free disialoside, we identified favourable glycan conformers to restrain modelling of the flexible α2,8-glycosidic linkage and were able to build the outward-facing, reducing-end Sia (Sia1) close to the e2 loop (Extended Data Fig. 21).

Binding of the ligand to the S1^A^ binding site is accompanied by local conformational changes, most conspicuously involving the displacement of the flanking e1 loop by 3 Å. W89 and T30 are brought in proximity to allow side chain hydrogen bonding, stabilising the *p1* pocket, while P33 shifts towards the *p2* pocket. Concomitantly, the N29 glycan, unresolved in the *apo* structure, becomes partially ordered and is displaced by 5 Å away from the S1^A^-S1^B^ interface (Supplementary Video 3). With the N-terminus stapled to the S1^A1^ core via a disulfide bond (C20-C156), the local changes in e1 are distally translated into long-range conformational changes. These extend all the way down to Y38, some 25 Å away from the binding pocket (Supplementary Video 4), located within a triple-strand hinge region that links the S1^A1^ and S1^A2^ subdomains (Fig. 4c, d). The resulting register shift between e1 segment (residues 29-37) and its neighbouring interacting partner (residues 73-81; indicated in black in Fig. 4c and 4d) seemingly drives the inward 9° rotation of the S1^A1^ subdomain about the S1^A1/A2^ axis (Fig. 4d, Supplementary Video 5).

### Molecular dynamics simulations independently predict S1^A^ conformational dynamics

The inherent flexibility of the disialoside binding pocket limits local resolution and the analysis of inter-residue interactions in our cryo-EM models. To gather atomistic insight into ligand binding, especially of Sia1, and the resulting shift in the protein conformational equilibrium, we performed MD simulations of the S1^A^ domain on an accumulated time scale of 70 μs.

Simulations starting from the ligand-bound cryo-EM *holo* structure revealed one dominating disialoside conformer in which the carboxylate of Sia1 interacts via a salt bridge with K84 while its 5-*N* is stabilised by a hydrogen bond with T82 (Fig. 5a, Extended Data Fig. 22-24, Supplementary Video 6).

Taking an unbiased MD approach into the conformational transition of e1, we used our structure of the *apo* S1^A^ domain as a starting model. The disialoside was placed into the binding pocket guided by the well-established orientation of 9-*O*-Ac-Sia2. Both the e1 and e2 loops showed pronounced dynamics in all trajectories as shown by a per-residue RMSD analysis (Extended Data Fig. 25-26). Saliently, conformational transitions observed in the e1 loop mirrored those identified upon comparison of the *apo* and closed *holo* cryo-EM models, even though the MD data were obtained fully independently (Fig. 5b, Extended Data Fig. 25, Supplementary Video 7). The observations were extended and corroborated by simulations with the S1^A^ domain of the HKU1-A N1 reference strain^28^, which differs from the Caen1 variant in that it carries a tyrosine instead of lysine at position 84 (Extended Data Fig. 26). All local conformational changes were observed, although a loss in stabilising interactions of Sia2 was noted, as would be expected due to the absence of K84 (Extended Data Fig. 27, Extended Data Tables 4-6).

In the *p1* pocket, two hydrogen bonds can form spontaneously, S86-L28 and T30-W89, with S86 and T30 orienting their hydrophilic hydroxyl groups away from the cavity. Alternatively, the crucial hydrogen bond with W89 can also be established with the neighbouring T31 sidechain (Extended Data Fig. 28). Flanking the *p2* pocket, interaction of P33 with F94 leads to a reduction in hydrophobic surface area and may contribute favourably to stability of the *holo* state of e1 in water. Further away from *p2*, long-range changes involving e1 residues R34 and S36 become apparent in the simultaneous breaking of two inter-strand backbone hydrogen bonds (S36-D76 and Y38-F74) and their re-formation with new partners (R34-D76 and S36-F74) in a ‘register shift’ motion (Fig. 5b), in full accordance with the observations by cryo-EM (Fig. 4c-d).

Two sets of control simulations of S1^A^ allowed us to infer a specific role of the ligand in the observed S1^A^ dynamics (Extended Data Fig. 26). In keeping with the inherent flexibility of the e1 loop, all individual e1 interactions can indeed also occur in the absence of the ligand. Without the ligand, however, these interactions remained highly dynamic. Yet, when the ligand encountered the alternative e1 state, either ‘naturally’ during the simulations or by simulations of a pre-built complex resembling the ‘*holo’* cryo-EM structure (Extended Data Fig. 26), this pattern changed substantially. The hallmark interactions, including the signature register shift in the S1^A1^-S1^A2^ hinge, reproducibly remained stable for several 100 ns. The collective results of cryo-EM and MD analyses indicate that ligand binding stabilises the shifted topology of the e1 element, apparently locking subdomain S1^A1^ in a state that allows subsequent conformational S1^B^ changes to occur.

## DISCUSSION

The dynamic sampling of open and closed conformations by sarbeco-and merbecovirus S proteins has become emblematic of how coronaviruses would balance host cell attachment and immune escape. The transition to the open state exposes subdomain S1^B^ for its binding to proteinaceous cell surface receptors and is also deemed crucial to allow protein refolding during S-mediated membrane fusion. Remarkably, however, with rare exception the pre-fusion S proteins from all other coronaviruses studied so far have all been observed in the closed state exclusively (Supplementary Table 1). Here we shed new light on this apparent contradiction by demonstrating that the S protein of a serotype A HKU1 strain can in fact transition into an open state, albeit not spontaneously but on a specific cue. Binding of the disialoside-based receptor 9-*O*-Ac-Sia(α2,8)Sia to S1^A^ triggers a major shift causing the S1^B^ subdomain to become exposed in a 1-up and eventually fully open, 3-up conformation. The exposure of S1^B2^ would allow for interactions with a putative secondary receptor and thus adds to the notion that such a receptor exists^16, 17^. Based on the collective data, we propose a model where binding to a primary sialoglycan-based receptor triggers opening of S1^B^, which in turn engages a yet unidentified secondary receptor required for entry (Fig. 6).

Four different S structures were identified that together capture a trajectory from a closed *apo* to a fully open *holo* conformation. The initial step, S1^A^ disialoside binding, converts the protein into a conformationally distinct state, still fully closed but primed for S1^B^ transition, transient yet stable enough to be detected in our analyses. The binding of the disialoside receptor analogue leads to various structural changes within the S1^A1^ subdomain. Most prominently, it stabilises an alternative topology of the e1 element, only fleetingly attained in the *apo* structure. Inward e1 displacement walls off one side of the 9-*O-*Ac-Sia binding site, deepening the *p1* pocket and adding to its hydrophobicity. Accommodation of the sialate-9-*O*-acetyl within *p1* may well act as the nucleating event from which other conformational changes follow. These extend to a distal hinge element which connects the S1^A1^ and S1^A2^ subdomains.

Although tempting to consider a causal mechanistic relationship between these conformational changes and S1^B^ transition, this would seem incongruous with observations of others. The topology of the e1 element in our type A HKU1 S *apo*-structure is atypical and differs from that in the S protein of type B HKU1 and those of betacoronavirus-1 variants OC43, bovine CoV and porcine hemagglutinating encephalomyelitis virus (Extended Data Fig. 29)^8,9, 24, 31^. In the *apo* structures of the latter proteins, the extended e1 element already adopts the topology of that in the type A HKU1 closed *holo* structure. Moreover, in the B-type HKU1 S *apo* structure, subdomains S1^A1^ and S1^A2^ are in similar spatial juxtaposition as in the A-type S *holo* conformation. Under the assumption that the other embecovirus S proteins also transition into an open conformation, they might do so via a distinct allosteric mechanism. We note, however, that cryo-EM models are based on averaging, and that it cannot be excluded that also in the HKU1-B and betacoronavirus-1 S proteins the e1 element continuously samples both topologies. If so, the transition of S1^B^ into the up position may critically depend on an increase in the lifetime of the shifted state as induced by S1^A^ ligand binding. Alternatively, the topology of the extended e1 element adopted in the type A HKU1 S *holo* structure may not be a trigger but rather a prerequisite to allow rotation of S1^A1^ around the S1^A1/A2^ axis and consequential remodelling of the S1^A^-S1^B^ interface—*i.e.* the phenomenon that seems most directly linked to S1^B^ expulsion. Thus, in A-type HKU1 S proteins, sialoside-dependent stabilisation of the e1 shift would be an additional precondition to be met to allow S1^B^ transition, *i.e.* on top of a generally conserved mechanism, the details of which remain to be defined. The difference between the A-and B-type spikes in their preferred *apo* topologies of the e1 element may have arisen from immune selection. Indeed, we recently demonstrated that the S1^A^ receptor binding site of OC43, which exhibits the shifted topology, is targeted by potent neutralising antibodies^15^.

The question remains why the transition into S1^B^ up conformations was not observed in our previous study of an OC43 S-receptor complex^9^. Possibly, the 9-*O*-Ac-Sia monosaccharide that was used as a receptor-analogue does not suffice to trigger the conformational changes and a more complex glycan may be required. Of note, OC43 S binds to α2,3 and α2,6-linked 9-*O*-Ac-Sias^9^, but displays a preference for 9-*O*-Ac-Sia(α2,8)Sia^10^. Evidence that OC43 S proteins can indeed transition to an open state with S1^B^ exposure comes from our recent observation of neutralising antibodies targeting cryptic S1^B^ epitopes. Moreover, virus neutralisation by these antibodies selected for resistance mutations in the e1 loop of S1^A^^15^. These results align with our present observations for HKU1, indicating that there is allosteric cross-talk between the S1^A^ and S1^B^ domains shared among embecoviruses. Hypervariable S1^A^ loop elements controlling both S1^B^ opening and S2’ proteolytic processing, as described for SARS-CoV-2, might even indicate that this is a universal feature of (beta)coronavirus S proteins^32, 33^. In this view, sarbeco-and merbecoviruses spontaneously exposing S1^B^ would not be exceptions but part of a mechanistic spectrum, with other CoVs, such as HKU1, relying on specific triggers such as binding to primary receptors via S1^A^. To our knowledge, this is the first description of a coronavirus S protein exposing its S1^B^ domain on cue. Our observations suggest that CoV attachment may be even more sophisticated than appreciated so far, with possibilities of dual receptor usage and priming of entry to escape immune detection.

## Methods

### Expression and purification of trimeric HKU1 S ectodomains

The sequence of a HKU1 type A spike (GenBank: ADN03339.1) encoding for the ectodomain (residues 12-1266) was cloned into the pCG2 expression vector with an exogenous CD5 signal peptide. At the 3’end, the coding sequence was ligated in frame with a GCN4 trimerization motif (IKRMKQIEDKIEEIESKQKKIENEIARIKKIK)^34, 35^, a thrombin cleavage site (LVPRGSLE), an 8-residue long Strep-Tag (WSHPQFEK) and a stop codon. The furin cleavage site at the S1/S2 junction was mutated from RRKRR to GGSGS to avert cleavage of the spike protein (Extended Data Fig. 30). The resulting construct was employed for transient expression in HEK293 cells and purified as previously described^36^. Briefly, after incubation of the cells for 5 days, spike glycoprotein was purified from cleared cell culture supernatants by affinity chromatography using StrepTactin beads (IBA) and eluted in 20 mM Tris-HCl, pH 8.0, 150 mM NaCl, 1 mM EDTA, 2.5 mM D-Biotin. The W89A mutant protein was produced as described previously^10^.

### Sample preparation for cryo-EM

For the *apo* complex, 3 µl of 4.3 µM HKU1 spike trimer was applied to QuantiFoil® R1.2/1.3 grids that had been glow-discharged for 30 seconds on a GloQube® (Quorum) at 20 mW power. The sample was applied at 4 °C and 95% relative humidity inside a Vitrobot Mark IV (Thermo Scientific). The grids were then blotted for 7 seconds with +2 blot force and plunge-frozen in liquid ethane. For the *holo* complex and W89A negative control, 7 µl of 4.3 μM WT or mutant HKU1 spike trimer was combined with 3 µl of 1 mM sugar, resulting in a final spike concentration of 3 μM and sugar concentration of 300 μM. The samples were then incubated at room temperature for ∼10 min prior to vitrification which was performed as described for the *apo* sample.

### Cryo-EM data acquisition

The *apo* and *holo* HKU1 spike samples were imaged on a Thermo Scientific™ Krios™ G4 Cryo-TEM equipped with a K3 direct electron detector and a BioContinuum® energy filter (Gatan) using EPU 2 acquisition software. The stage was pre-tilted to 30° to improve the orientation distribution of the particles. A total of 4207 movies for *apo* spike and 4,065 movies for the *holo* spike were collected at a super-resolution pixel size of 0.415 Å/pixel, with 40 fractions per movie and a total dose of 46 e^-^/Å^2^. Defocus targets cycled from -1.5 to -2.5 microns.

The W89A mutant HKU1 spike incubated with disialoside was imaged on a Thermo Scientific™ Glacios Cryo-TEM equipped with a Falcon 4 direct electron detector using EPU 2 acquisition software. The stage was pre-tilted to 30° to improve the orientation distribution of the particles. A total of 896 movies were collected at 0.92 Å/pixel with 40 fractions per movie and a total dose of 42 e^-^/Å^2^. Defocus targets cycled from -1.5 to -2.5 microns. A summary of all data collection parameters is shown in supplementary table 2.

### Single-particle image processing

For the *apo* complex, patch motion correction, using an output F-crop factor of 0.5, and patch CTF estimation were performed in cryoSPARC live^37^. Micrographs with a CTF estimated resolution of worse than 10 Å were discarded, leaving 4202 images for further processing. The blob picker tool was then used to select 9144772 particles which were then extracted in a 100-pixel box (Fourier binned 4 × 4) and then exported to cryoSPARC for further processing. A single round of 2D classification was performed, after which 183886 particles were retained. Ab initio reconstruction generated one well-defined reconstruction of the closed HKU1 spike. Particles belonging to this class were then re-extracted in a 300-pixel box. During extraction, particles were Fourier binned by a non-integer value, resulting in a final pixel size of 1.1067 Å. Subsequently, non-uniform refinement was performed on the extracted particles with C3 symmetry imposed^38^, yielding a reconstruction with a global resolution of 3.3 Å. Subsequently, each particle from the C3 symmetry–imposed reconstruction was assigned three orientations corresponding to its symmetry-related views using the symmetry expansion job. A soft mask encompassing one S1^A^ domain was made in UCSF Chimera^39^, and used for local refinement of the expanded particles, yielding a map with a global resolution of 3.8 Å.

For the *holo* complex, patch motion correction, using an output F-crop factor of 0.5, and patch CTF estimation were performed in cryoSPARC live^37^. Micrographs with a CTF estimated resolution of worse than 10 Å were discarded, leaving 4,045 images for further processing. The blob picker tool was then used to select 956,697 particles which were then extracted in a 100-pixel box (Fourier binned 4 × 4) and then exported to cryoSPARC for further processing. Four parallel rounds of 2D classification were performed, using an initial classification uncertainty value of 1, 2, 4 or 6. Subsequently, the well-defined spike classes were selected from each 2D run and combined. Duplicate particles were then removed, after which 169,728 particles were retained. Ab initio reconstruction generated two classes corresponding to the closed and 3-up spike trimer. Particles from these two classes were used as the input for a second round of ab initio reconstruction which produced two classes corresponding to the 3-up and 1-up spike trimer, although the latter appear to be a convolution of 1-up and closed particles. These two volumes were then used as initial models for a round of heterogenous refinement. To avoid missing spike particles which may have been removed during initial stringent selection of 2D classes, heterogenous refinement was performed on a larger particle stack of 895,888 particles, from which only carbon classes had been removed from the initial stack. Heterogenous refinement produced two well-defined reconstructions of the 3-up and 1-up conformations. Particles corresponding to the 3-up class were subjected to a single round of 2D classification and the clearly defined spike classes were selected. These were then re-extracted in a 300-pixel box. During extraction, particles were Fourier binned by a non-integer value, resulting in a final pixel size of 1.1067 Å. Subsequently, non-uniform refinement was performed on the extracted particles with C3 symmetry imposed^38^, yielding a reconstruction with a global resolution of 3.7 Å. Because of the apparent heterogeneity in the 1-up sample, an additional round of heterogenous refinement was performed on the 895,888-particle stack, using higher quality initial models, namely the fully refined 3-up map and the 1-up map obtained from the second round of ab initio reconstruction. Heterogenous refinement produced well-defined reconstructions of the 3-up and 1-up conformations. Particles corresponding to both classes were individually subjected to a single round of 2D classification and the clearly defined spike classes were selected. These were then individually re-extracted in a 300-pixel box. During extraction, particles were Fourier binned by a non-integer value, resulting in a final pixel size of 1.1067 Å. Subsequently, non-uniform refinement was performed on the extracted particles with C3 or C1 symmetry imposed, yielding reconstructions with global resolutions of 3.56 and 4.13 Å for the 3-up and 1-up conformations, respectively. After global refinement, a soft mask encompassing one S1^A^ domain of the 3-up sample was made in UCSF Chimera. Local refinement was then performed on the 3-up particles, yielding a map with a global resolution of 4.19 Å. The particles belonging to the 1-up reconstruction were subjected to another round of heterogenous refinement, which produced two clear reconstructions of the closed and 1-up spike. Non-uniform refinement was performed on both sets of particles with C3 or C1 symmetry imposed, yielding reconstructions with global resolutions of 3.68 and 4.68 Å for the closed and 1-up conformations, respectively. For the closed spike, each particle from the C3 symmetry–imposed reconstruction was assigned three orientations corresponding to its symmetry-related views using the symmetry expansion job. A soft mask encompassing one S1^A^ domain was made in UCSF Chimera^39^, and the symmetry-expanded particles were subjected to masked 3D variability analysis^40^. Local refinement was then performed on the particles belonging to the best resolved cluster, yielding a map with a global resolution of 4.13 Å.

For the W89A mutant HKU1 spike incubated with disialoside, patch motion correction was performed in MotionCor2^41^, implemented through Relion version 3.1.1^42^. The motion corrected micrographs were then imported into cryoSPARC for patch CTF estimation and further processing steps^37^. The blob picker tool was used to select 215843 particles which were then extracted in a 100-pixel box (Fourier binned 4 × 4). A single round of 2D classification was performed, after which 38838 particles were retained. Ab initio reconstruction generated one well-defined reconstruction of the closed HKU1 spike. Particles belonging to this class were then re-extracted in a 300-pixel box. During extraction, particles were Fourier binned by a non-integer value, resulting in a final pixel size of 1.2267 Å. Subsequently, non-uniform refinement was then performed on the extracted particles with C3 symmetry imposed^38^, yielding a reconstruction with a global resolution of 5.1 Å. Subsequently, each particle from the C3 symmetry–imposed reconstruction was assigned three orientations corresponding to its symmetry-related views using the symmetry expansion job. A soft mask encompassing one S1^A^ domain was then made in UCSF Chimera^39^, and used for local refinement of the expanded particles, yielding a map with a global resolution of 5.4 Å.

The “Gold Standard” Fourier shell correlation (FSC) criterion (FSC = 0.143) was used for calculating all resolution estimates, and 3D-FSC plots were generated in cryoSPARC^43^. To facilitate model building, globally refined maps were sharpened using DeepEMhancer (version 0.13)^44^, as implemented in COSMIC2 (ref. 54)^45^, or filtered by local resolution in cryoSPARC.

### Modelling

Initially, a Phyre2^46^ generated homology model (template pdb 6NZK^17^; the embecovirus OC43 S) for HKU1-A S was rigid body fitted into the *apo* state cryo-EM map using UCSF Chimera^39^ “Fit in map”. The crystal structure of HKU1-A S1^B^ (pdb 5KWB,^17^) was used to replace the equivalent S1^B^ domain in the homology model due to clearly wrong homology modelling.

Models were refined by performing iterative cycles of manual model building using Coot^47^ and real space refinement using Phenix^48^. The Coot carbohydrate module^49^ was used for building N-linked glycans, which were manually inspected and corrected. The *apo* state was modelled first, due to its highest resolution. Subsequently, the closed *holo*, the *holo* 3-up and the *holo* 1-up were modelled in that order, using previous models as a starting point. For the initial *holo* (closed) S1^A^ model, Namdinator^50^ was used for flexible fitting in a locally refined and unsharpened map for the closed *holo* S1^A^. Model validation was performed using Molprobity^51^.

Elbow^52^ was used to generate ligand restraints for the 9-*O*-acetylated terminal sialic acid based on the “MJJ” ligand in the OC43 S cryo-EM structure (PDB 6NZK^9^), after which atom names were manually modified to be consistent with the earlier standard MJJ model and general sialic acid atom numbering, and the O2-attached methyl linker atoms of the original MJJ ligand were trimmed. Since there is no standard MJJ-SIA α2,8 linkage defined in currently used software packages, we used MD-based restraints (see below) to model this glycosidic linkage of the disialoside. The following restraints were used for the glycosidic linkage between the terminal 9-*O* acetylated sialic acid (ligand code MJJ) and the penultimate sialic acid (ligand code SIA) based on the most common solution conformer: bond distance C2-O8 of 1.38 Å (σ of 0.01 Å); bond angles of 109.5° for O8-C2-O6 and for O8-C2-C3, 114.5° for O8-C2-C1 (all σ of 2.0°); dihedral angles of 295.0° for C1-C2-O8-C8 and of 122° for C2-O8-C8-C7 (both σ of 5.0°).

### Molecular dynamics simulations

Starting structures of the molecular systems were built based on the cryo-EM structures of HKU1 (this work) using the graphical interface of YASARA^53^. The N-glycans were attached to the protein based on data from quantitative site-specific N-linked analysis of HKU1 S^29^. Models of the complexes with α-Neu5Ac-(2-8)-α-Neu5Ac-OMe were build based on the *holo* and *apo* versions of S1^A^ (residues 14-299). The ligand was positioned manually into the binding site guided by interactions found in PDB entry 6NZK (hCoV-OC43). The HKU1 N1 sequence was taken from GenBank entry NC_006577.2.

In general, the systems were solvated in 0.9% NaCl solution (0.15 M) and simulations were performed at 310 K using periodic boundary conditions using the AMBER14 force field^54–56^. The box size was rescaled dynamically to maintain a water density of 0.996 g/ml. Simulations were performed using YASARA with GPU acceleration in ‘fast mode’ (4 fs time step)^57^ on ‘standard computing boxes’ equipped *e.g.* with one 12-core i9 CPU and NVIDIA GeForce GTX 1080 Ti.

The fully glycosylated ectodomain system (590814 atoms) was simulated for 250 ns with a performance of about 4 ns/day. Molecular systems based on S1^A^ only were smaller (approx. 32500-56200 atoms, depending on the size of the N-glycans attached) and were sampled for an accumulated timescale of approx. 20 µs for the Caen1 sequence (*apo* + disialoside ligand, 5 µs, 6 simulations; *holo*, 15 µs, 27 simulations) and 52 µs for the N1 sequence (*apo*, 12 µs, 13 simulations; *apo* + disialoside ligand, 23 µs, 34 simulations; *holo*, 17 µs, 22 simulations) with individual simulations reaching up to 1.6 µs. The performance was about 100-200 ns/day. Distances shown in Fig. 5b were calculated from an example trajectory (Extended Data Fig. 25) between the following atoms: L28:O-S86:OG, P33:CG-F94:CA, T30:O-W89:NE1, S36:N-F74:O, R34:O-D76:N. Conformational Analysis Tools (CAT, http://www.md-simulations.de/CAT/) was used for analysis of trajectory data, general data processing and generation of scientific plots. VMD^58^ was used to generate molecular graphics.

### Analysis and visualization

S interface areas were calculated using PDBePISA^59^. Surface colouring of HKU1-A S according to sequence conservation was performed using Consurf^60^ and visualised in UCSF ChimeraX ^61^. The UCSF Chimera “MatchMaker” tool was used to obtain root mean square deviation values, using default settings. Domain rotations were calculated with CCP4^62^ Superpose ^63^. Figures were generated using UCSF ChimeraX^61^ and biorender.com. Structural biology applications used in this project were compiled and configured by SBGrid^64^.

## Data availability

The atomic models of the *apo*, *holo*, 1-up and 3-up HCoV-HKU1 spike have been deposited to the Protein Data Bank under the accession codes 8OHN, 8OPM, 8OPN and 8O9-OPO. The globally and locally refined cryo-EM maps have been deposited to the Electron Microscopy Data Bank under the accession codes EMD-16882, EMD-17076, EMD-17077, EMD-17078, EMD-17079, EMD-17080, EMD-17081, EMD-17082 and EMD-17083.

## Acknowledgments

We thank Ronald Dijkman for providing the HKU1 Caen1 sequence and Jolanda de Groot-Mijnes for critically reading the manuscript. We are grateful for computer time provided by BIOGNOS AB, Göteborg. This work was supported by the China Scholarship Council 2014-03250042 (YL). This work made use of the Dutch national e-infrastructure with the support of the SURF Cooperative using grant no. EINF-2453, awarded to DLH. RC acknowledges funding by the Deutsche Forschungsgemeinschaft (494746248). RJGH is funded by a Dutch research council NWO-XS grant (OCENW.XS22.3.110). GJB is supported by an ERC Advanced Grant (SWEETPROMISE, 101020769), MFP and DLH by NWO Veni grants (VI.Veni.202.271 and VI.Veni.212.102, respectively). JS is funded by the Dutch Research Council NWO Gravitation 2013 BOO, Institute for Chemical Immunology (ICI; 024.002.009).

## Author contributions

YL, RJdG and DLH conceived the project; YL, MF and DLH designed the experiments; YL designed and cloned the protein constructs and carried out protein expression and purification; ID, ZL, FJMvK, JS, B-JB, G-JB and MF provided access to equipment and reagents; ID performed cryo-EM sample preparation and data collection; DLH processed the cryo-EM data; MFP and DLH built and refined the atomic models. MF carried our molecular dynamics simulations; MFP, RC, MF and DLH analysed and visualised the data; MFP, MF and DLH curated the data. RJdG and DLH supervised the project. MFP, RC, RJGH, RJdG and DLH carried out project administration. MFP, YL, RJGH, RJdG and DLH obtained funding. MFP, RC, RJGH, RJdG and DLH wrote the first draft of the manuscript. All authors contributed to reviewing and editing subsequent versions.

