## Supplementary Information for "Sialoglycan binding triggers spike opening in a human coronavirus"

**This PDF file includes:**

Figs. S1 to S30
Tables S1 to S6
Captions for Videos S1 to S7

**Other Supplementary Materials for this manuscript include the following:**

Videos S1 to S7

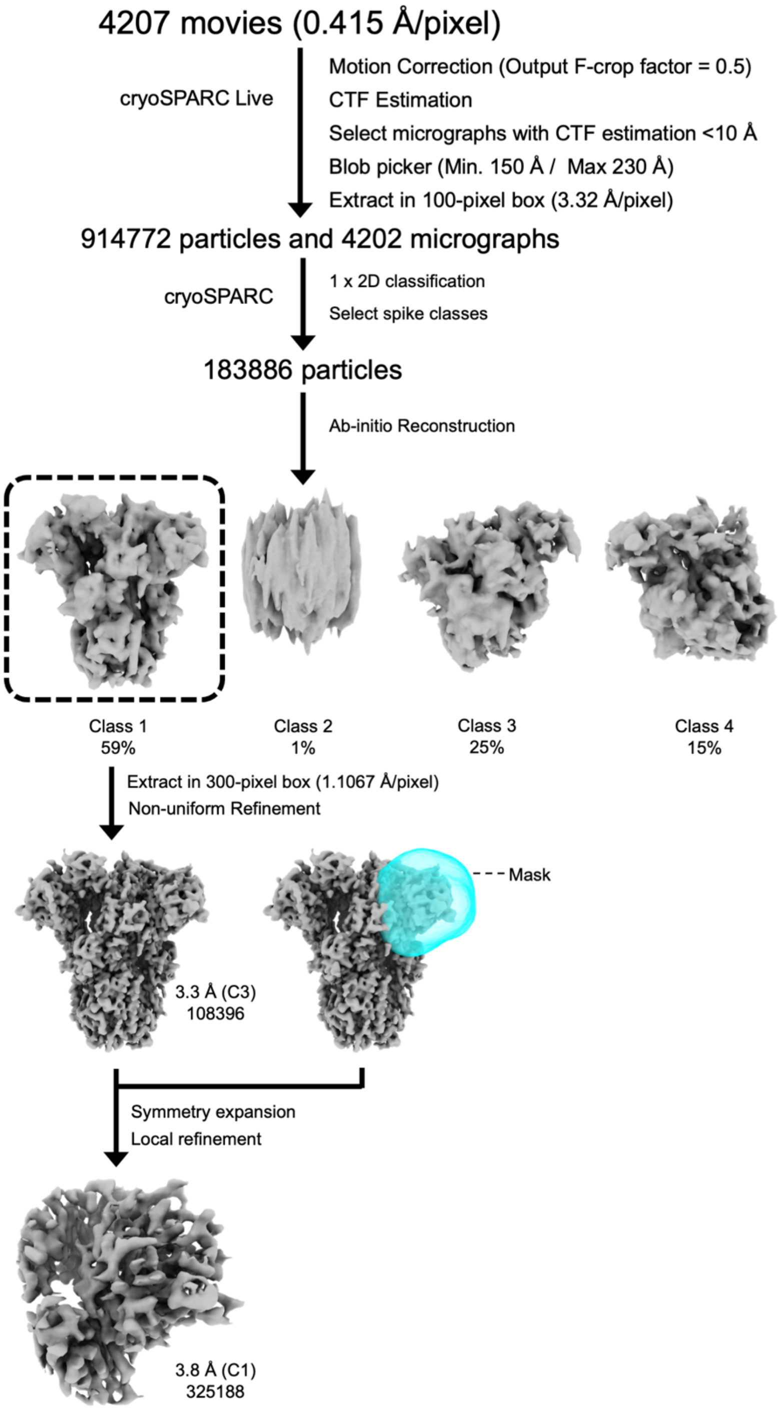

**Extended Data Fig. 1. Cryo-EM data processing pipeline for the *apo* HKU1-A spike glycoprotein.**

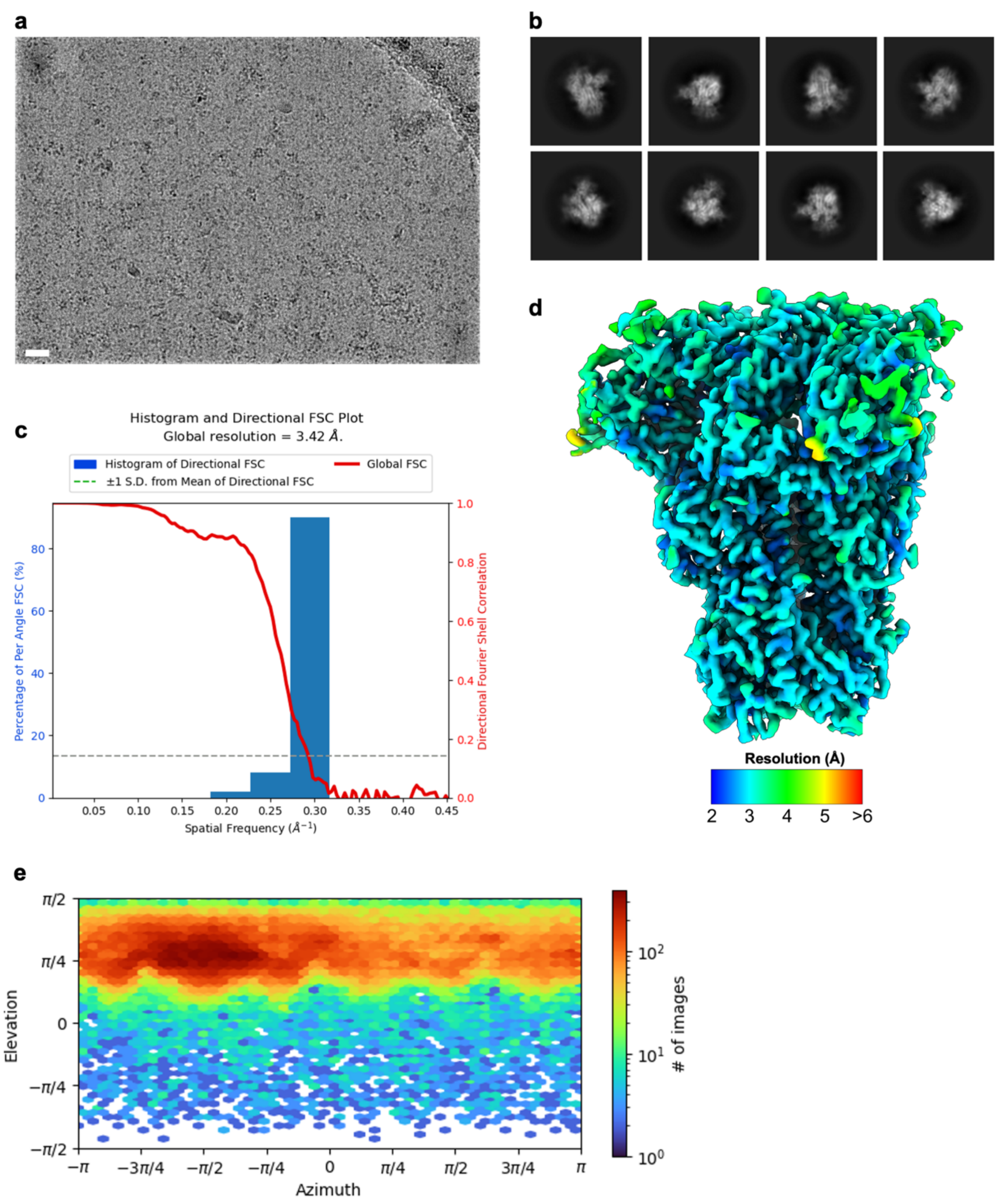

**Extended Data Fig. 2. Cryo-EM data processing of the *apo* HKU1-A S ectodomain.**

**a**, Representative motion-corrected micrograph of the *apo* HKU1-A spike ectodomains embedded in vitreous ice. Scale bar = 50 nm. **b**, Representative reference-free 2D class averages generated in cryoSPARC. **c**, 3DFSC plot for the 3.4 Å resolution globally refined reconstruction. **d**, DeepEMhancer filtered EM density map for the *apo* HKU1-A spike ectodomains coloured according to local resolution which was calculated in cryoSPARC. **e**, Angular distribution plot calculated in cryoSPARC for particle projections in the globally refined map.

```
HKU1-A      MLLIIFILPTTLAVIGDFNCTNFAINDLNTTIPRISEYVVDVSYGLGTYYILDRVYLNTT 60
HKU1-B      MFLIIFILPTTLAVIGDFNCTNSFINDYNKTIPRISEDVVDVSLGLGTYYVLNRVYLNTT 60
            *:********************  *** *.******* ***** ******:*:*******

HKU1-A      ILFTGYFPKSGANFRDLSLKGTTKLSTLWYQKPFLSDFNNGIFSRVKNTKLYVNKTLYSE 120
HKU1-B      LLFTGYFPKSGANFRDLALKGSIYLSTLWYKPPFLSDFNNGIFSKVKNTKLYVNNTLYSE 120
            :****************:***:  ******: ************:*********:*****

HKU1-A      FSTIVIGSVFINNSYTIVVQPHNGVLEITACQYTMCEYPHTICKSIGSSRNESWHFDKSE 180
HKU1-B      FSTIVIGSVFVNTSYTIVVQPHNGILEITACQYTMCEYPHTVCKSKGSIRNESWHIDSSE 180
            **********:*.***********:****************:*** ** ******:*.**

HKU1-A      PLCLFKKNFTYNVSTDWLYFHFYQERGTFYAYYADSGMPTTFLFSLYLGTLLSHYYVLPL 240
HKU1-B      PLCLFKKNFTYNVSADWLYFHFYQERGVFYAYYADVGMPTTFLFSLYLGTILSHYYVMPL 240
            **************:************.******* **************:******:**

HKU1-A      TCNAISSNTDNETLQYWVTPLSKRQYLLKFDDRGVITNAVDCSSSFFSEIQCKTKSLLPN 300
HKU1-B      TCNAISSNTDNETLEYWVTPLSRRQYLLNFDEHGVITNAVDCSSSFLSEIQCKTQSFAPN 300
            **************:*******:*****:**::*************:*******:*: **

HKU1-A      TGVYDLSGFTVKPVATVHRRIPDLPDCDIDKWLNNFNVPSPLNWERKIFSNCNFNLSTLL 360
HKU1-B      TGVYDLSGFTVKPVATVYRRIPNLPDCDIDNWLNNVSVPSPLNWERRIFSNCNFNLSTLL 360
            *****************:****:*******:****..*********:*************

HKU1-A      RLVHTDSFSCNNFDESKIYGSCFKSIVLDKFAIPNSRRSDLQLGSSGFLQSSNYKIDTTS 420
HKU1-B      RLVHVDSFSCNNLDKSKIFGSCFNSITVDKFAIPNRRRDDLQLGSSGFLQSSNYKIDISS 420
            ****.*******:*:***:****:**.:******* **.****************** :*

HKU1-A      SSCQLYYSLPAINVTINNYNPSSWNRRYGFNNFNLSSHSVVYSRYCFSVNNTFCPCAKPS 480
HKU1-B      SSCQLYYSLPLVNVTINNFNPSSWNRRYGFGSFNLSSYDVVYSDHCFSVNSDFCPCADPS 480
            ********** :******:***********..*****:.**** :*****. *****.**

HKU1-A      FASSCKSHKPPSASCPIGTNYRSCESTTVLDHTDWCRCSCLPDPITAYDPRSCSQKKSLV 540
HKU1-B      VVNSCAKSKPPSAICPAGTKYRHCDLDTTLYVKNWCRCSCLPDPISTYSPNTCPQKKVVV 540
            ...** . ***** ** **:** *:  *.*  .:***********::*.*.:* *** :*

HKU1-A      GVGEHCAGFGVDEEKCGVLDGSYNVSCLCSTDAFLGWSYDTCVSNNRCNIFSNFILNGIN 600
HKU1-B      GIGEHCPGLGINEEKCGTQL--NHSSCFCSPDAFLGWSFDSCISNNRCNIFSNFIFNGIN 598
            *:**** *:*::*****.     : **:** *******:*:*:************:****

HKU1-A      SGTTCSNDLLQPNTEVFTDVCVDYDLYGITGQGIFKEVSAVYYNSWQNLLYDFNGNIIGF 660
HKU1-B      SGTTCSNDLLYSNTEISTGVCVNYDLYGITGQGIFKEVSAAYYNNWQNLLYDSNGNIIGF 658
            **********  ***: *.***:*****************.***.******* *******

HKU1-A      KDFVTNKTYNIFPCYAGRVSAAFHQNASSLALLYRNLKCSYVLNNISLATQP-YFDSYLG 719
HKU1-B      KDFLTNKTYTILPCYSGRVSAAFYQNSSSPALLYRNLKCSYVLNNISFISQPFYFDSYLG 718
            ***:*****.*:***:*******:**:** *****************: :** *******

HKU1-A      CVFNADNLTDYSVSSCALRMGSGFCVDYNSPSSSSSRRKRRSISASYRFVTFEPFNVSFV 779
HKU1 B      CVLNAVNLTSYSVSSCDLRMGSGFCIDYALPS---SRRKRRGISSPYRFVTFEPFNVSFV 775
            **:** ***.****** ********:**  **   ******.**: **************

HKU1-A      NDSIESVGGLYEIKIPTNFTIVGQEEFIQTNSPKVTIDCSLFVCSNYAACHDLLSEYGTF 839
HKU1-B      NDSVETVGGLFEIQIPTNFTIAGHEEFIQTSSPKVTIDCSAFVCSNYAACHDLLSEYGTF 835
            ***:*:****:**:*******.*:******.********* *******************

HKU1-A      CDNINSILDEVNGLLDTTQLHVADTLMQGVTLSSNLNTNLHFDVDNINFKSLVGCLGPHC 899
HKU1-B      CDNINSILNEVNDLLDITQLQVANALMQGVTLSSNLNTNLHSDVDNIDFKSLLGCLGSQC 895
            ********:***.*** ***:**::**************** *****:****:**** :*

HKU1-A      GSSSRSFFEDLLFDKVKLSDVGFVEAYNNCTGGSEIRDLLCVQSFNGIKVLPPILSESQI 959
HKU1-B      GSSSRSLLEDLLFNKVKLSDVGFVEAYNNCTGGSEIRDLLCVQSFNGIKVLPPILSETQI 955
            ******::*****:*******************************************:**
```

```
HKU1-A      SGYTTAATVAAMFPPWSAAAGIPFSLNVQYRINGLGVTMDVLNKNQKLIATAFNNALLSI 1019
HKU1-B      SGYTTAATVAAMFPPWSAAAGVPFSLNVQYRINGLGVTMDVLNKNQKLIANAFNKALLSI 1015
            *********************:****************************.***:*****

HKU1-A      QNGFSATNSALAKIQSVVNSNAQALNSLLQQLFNKFGAISSSLQEILSRLDALEAQVQID 1079
HKU1-B      QNGFTATNSALAKIQSVVNANAQALNSLLQQLFNKFGAISSSLQEILSRLDNLEAQVQID 1075
            ****:**************:******************************* ********

HKU1-A      RLINGRLTALNAYVSQQLSDISLVKLGAALAMEKVNECVKSQSPRINFCGNGNHILSLVQ 1139
HKU1-B      RLINGRLTALNAYVSQQLSDITLIKAGASRAIEKVNECVKSQSPRINFCGNGNHILSLVQ 1135
            *********************:*:* **: *:****************************

HKU1-A      NAPYGLLFMHFSYKPISFKTVLVSPGLCISGDVGIAPKQGYFIKHNDHWMFTGSSYYYPE 1199
HKU1-B      NAPYGLLFIHFSYKPTSFKTVLVSPGLCLSGDRGIAPKQGYFIKQNDSWMFTGSSYYYPE 1195
            ********:****** ************:*** ***********:** ************

HKU1-A      PISDKNVVFMNTCSVNFTKAPLVYLNHSVPKLSDFESELSHWFKNQTSIAPNLTLNLHTI 1259
HKU1-B      PISDKNVVFMNSCSVNFTKAPFIYLNNSIPNLSDFEAELSLWFKNHTSIAPNLTFNSH-I 1254
            ***********:*********::***:*:*:*****:*** ****:********:* * *

HKU1-A      NATFLDLYYEMNLIQESIKSLNNSYINLKDIGTYEMYVKWPWYVWLLISFSFIIFLVLLF 1319
HKU1-B      NATFLDLYYEMNVIQESIKSLNSSFINLKEIGTYEMYVKWPWYIWLLIVILFIIFLMILF 1314
            ************:*********.*:****:*************:**** : *****::**

HKU1-A      FICCCTGCGSACFSKCHNCCDEYGGHHDFVIKTSHDD                        1356
HKU1-B      FICCCTGCGSACFSKCHNCCDEYGGHNDFVIKASHDD                        1351
            **************************:*****:****
```

**Extended Data Fig. 3. Sequence alignment of HKU1-A and HKU1-B spike glycoproteins.**

Comparison of HKU1 S protein sequences from genotypes A (Caen1, GenBank entry ADN03339.1) and B (isolate N5, NCBI entry Q0ZME7) aligned using the ClustalW algorithm[1].

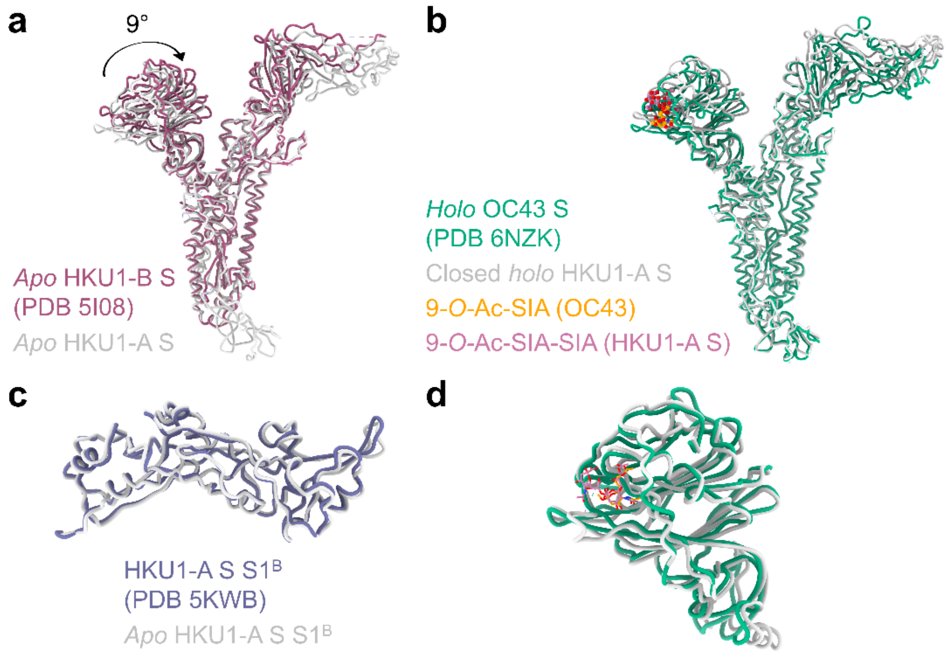

**Extended Data Fig. 4. Comparisons of our HKU1-A S structures with previously published embecovirus spike structures**

**a**, Comparison of our HKU1-A S structures (in grey) with the previously published structure of full-length HKU1-B S (dark pink). **b**, Comparison of a protomer of the OC43 S (green) bound with a 9-*O*-acetylated sialic acid (orange) with our closed *holo* $S1^A$. **c**, Comparison of our HKU1-A $S1^B$ domain structure with the previously published HKU1-A $S1^B$ domain crystal structure (purple). **d**, Close up comparison of the $S1^A$ domains of OC43 and our HKU1-A aligning on the $S1^A$ domain instead of the whole spike (same colouring as panel **b**).

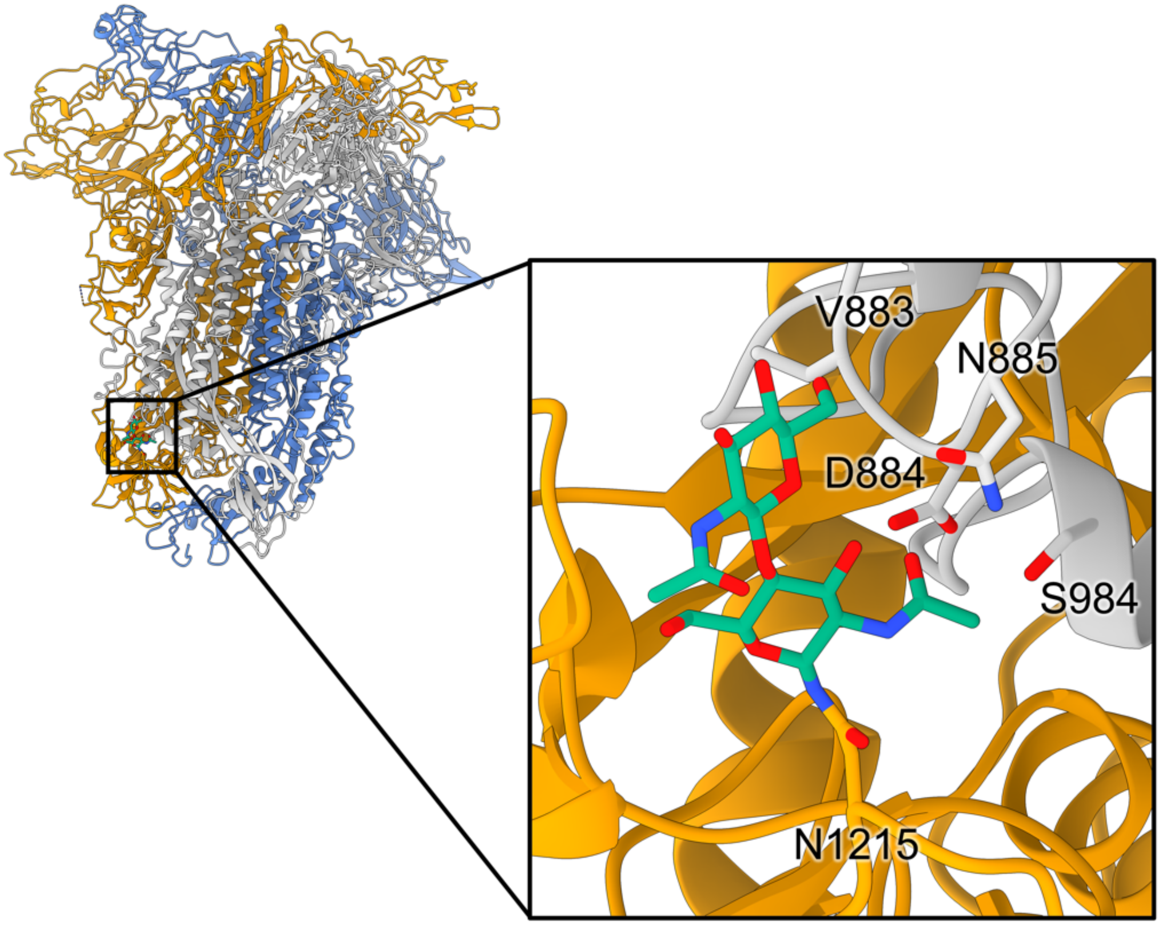

**Extended Data Fig. 5. N-linked glycosylation of N1215 stabilises the trimer at its base via contacts with the counter-clockwise neighbouring protomer.**

Residues from the neighbouring protomer (in grey) contacting this N-linked glycan (green) are indicated as sticks.

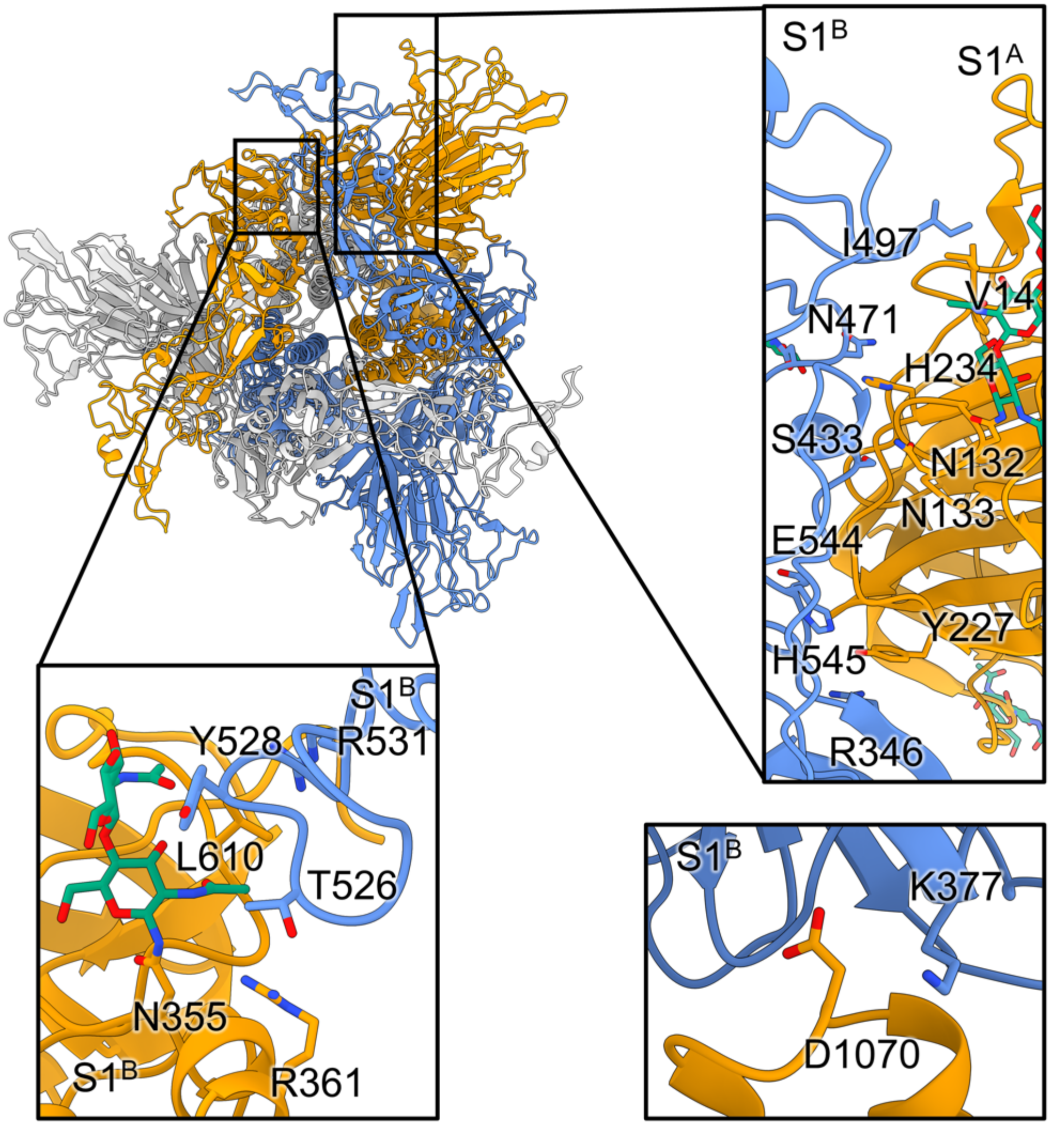

**Extended Data Fig. 6. Several minor interfaces stabilise the S1$^{B}$ domains in a downward orientation.**

Several minor interfaces stabilise the S1$^{B}$ domains in a downward orientation in the *apo* state, such as S1$^{B}$-S1$^{B}$ (bottom left), S1$^{B}$-S1$^{A}$ (top right) and S1$^{B}$-S2 (bottom right). The S1$^{B}$-S1$^{B}$ interface (bottom left) is stabilised by an N-linked glycan on N355.

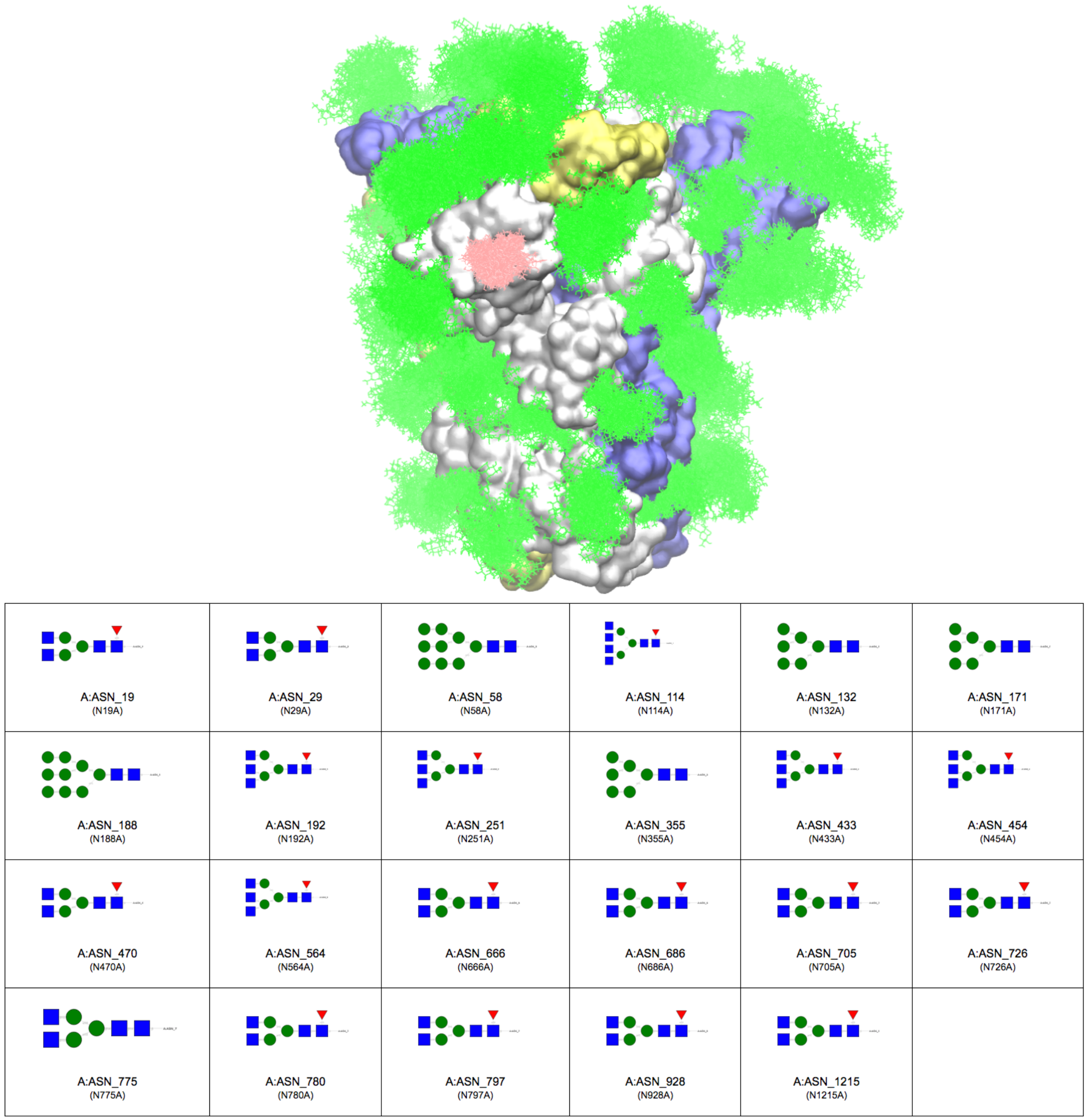

**Extended Data Fig. 7. Glycan coverage of the HKU1-A spike ectodomain.**

MD-derived glycan coverage map of the HKU1 S ectodomain (250 ns, 310 K). Full N-glycans (as shown for chain A) were attached based on previously published data[2] where available. The spike protomers are coloured grey, blue and yellow and the N-linked glycans and bound disialoside are coloured green and pink, respectively. To highlight the dynamics of the N-glycans, 250 snapshots extracted at time intervals of 1 ns are shown overlayed using VMD. The table with the SNFG representations of the N-glycans was generated using Conformational Analysis Tools (CAT, http://www.md-simulations.de/CAT/).

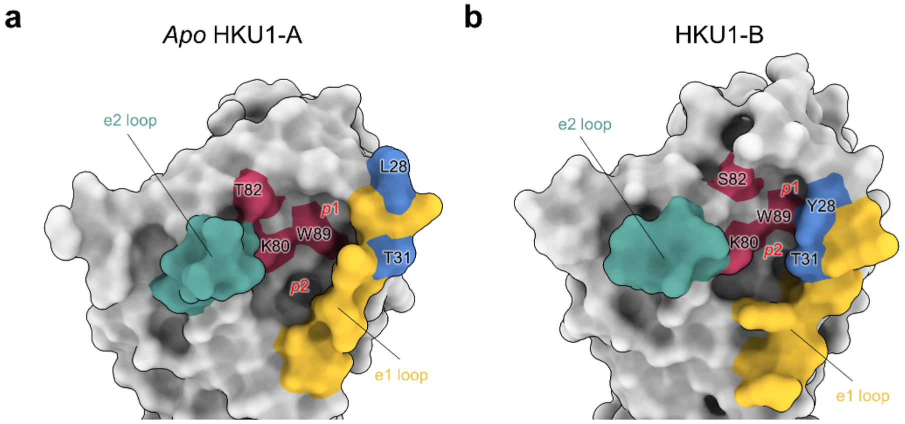

**c**

```
HKU1-A      VIGDFNCTNFAINDLNTTVPRISEYVVDVSYGLGTYYILDRVYLNTTILFTGYFPKSGAN
HKU1-B      VIGDFNCTNSFINDYNKTIPRISEDVVDVSLGLGTYYVLNRVYLNTTLLFTGYFPKSGAN
            *********  *** *.*:***** ***** ******:*:*******:************

HKU1-A      FRDLSLKGTTYLSTLWYQKPFLSDFNNGIFSRVKNTKLYVNKTLYSEFSTIVIGSVFINN
HKU1-B      FRDLALKGSIYLSTLWYKPPFLSDFNNGIFSKVKNTKLYVNNTLYSEFSTIVIGSVFVNT
            ****:***: *******: ************:*********:***************:*.
```

**Extended Data Fig. 8. Comparison of the *apo* HKU1-A and HKU1-B sialic acid binding sites**
**a**, Surface representation of the *apo* HKU1-A sialic acid binding site. Residues critical for sialic acid binding are coloured ruby and selected e1 loop residues are coloured blue. The location of the *p*1 and *p*2 pockets are indicated. **b**, Surface representation of the HKU1-B sialic acid binding site (PDB ID: 5I08)[3], same colouring as panel **a**. **c**, alignment of HKU1-A and -B S1$^{A}$ segments involved in sialic acid binding. Important residues highlighted in panel **a** and **b** are again highlighted in red and blue.

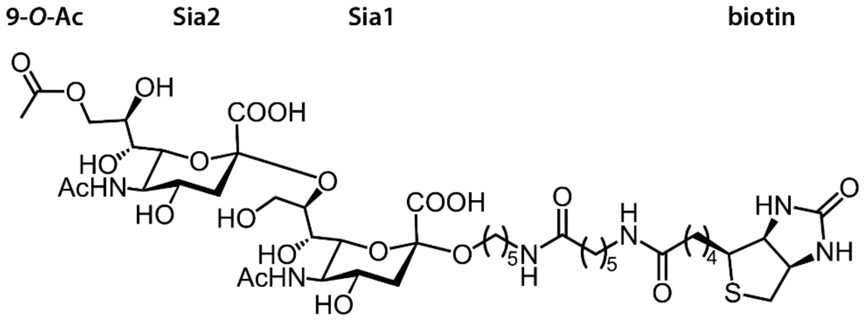

**Extended Data Fig. 9. Structure of the receptor analogue 9-*O*-Ac-Neu5Ac-α2,8-Neu5Ac-Lc-biotin**

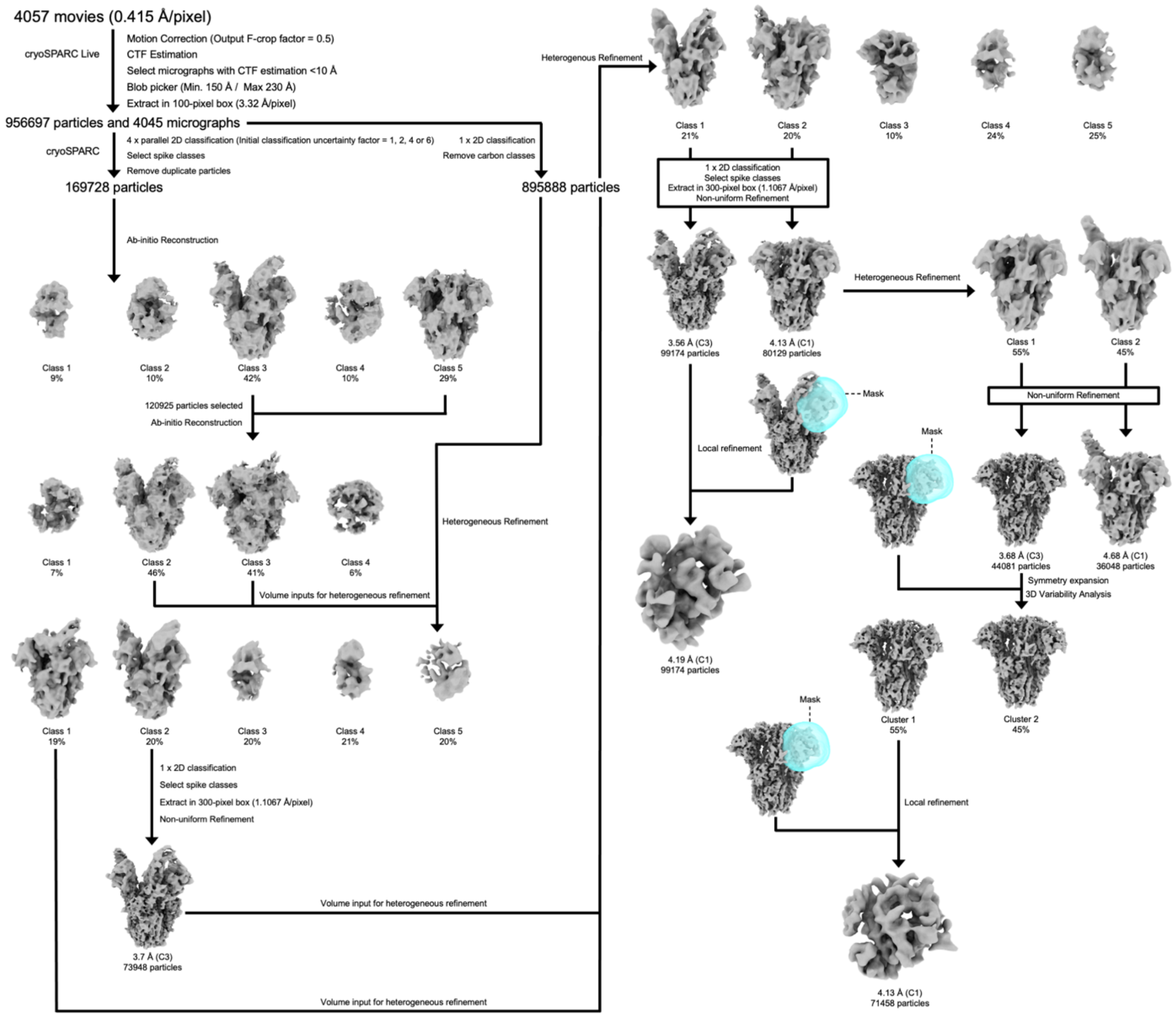

**Extended Data Fig. 10. Cryo-EM data processing pipeline for the *holo* HKU1-A spike glycoprotein.**

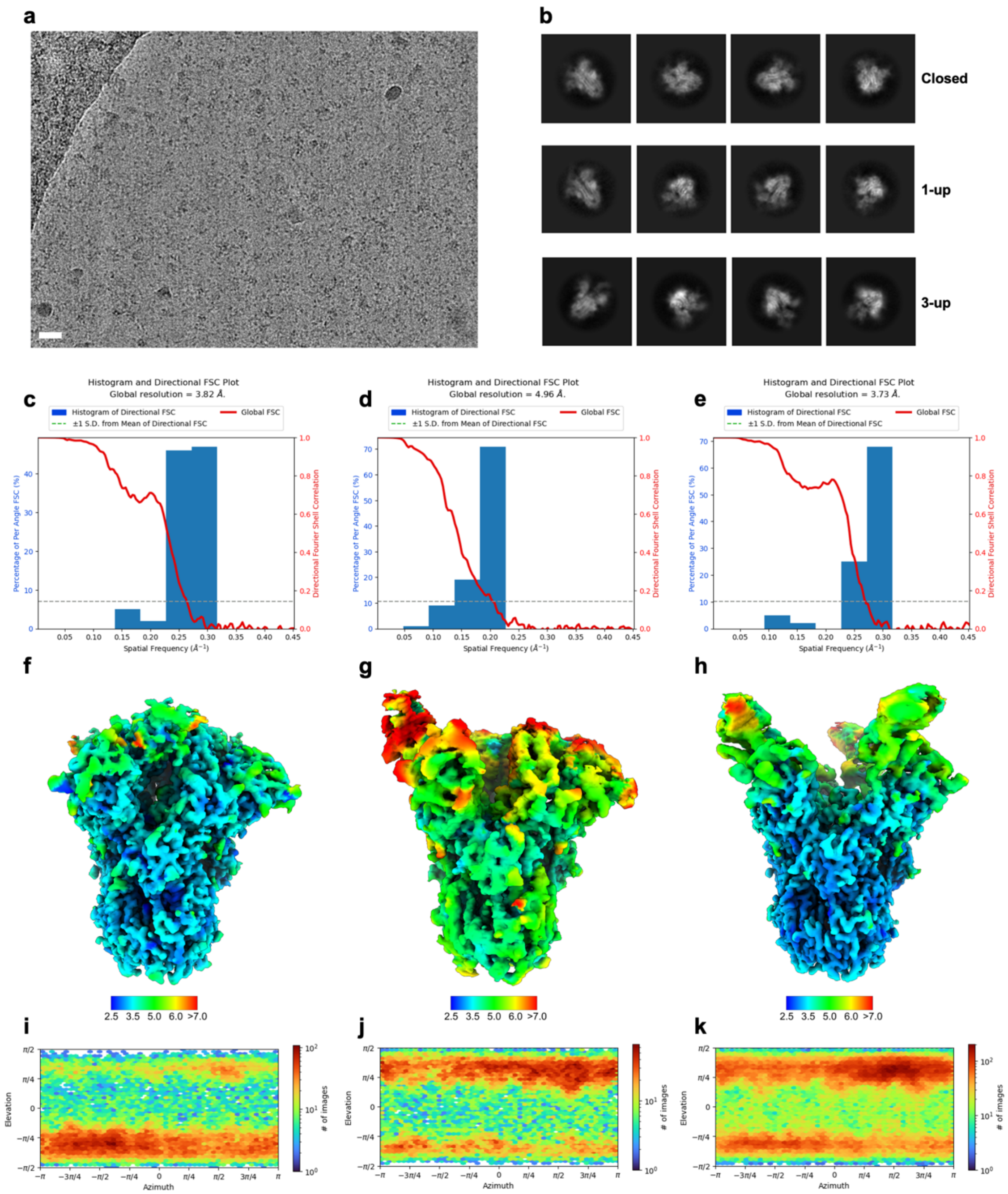

**Extended Data Fig. 11. Cryo-EM data processing of the *holo* HKU1-A S ectodomain**

**a**, Representative motion-corrected micrograph of the disialoside-incubated HKU1-A spike ectodomains embedded in vitreous ice. Scale bar = 50 nm. **b**, Representative reference-free 2D class averages of the closed, 1-up and 3-up reconstructions generated in cryoSPARC. **c**, 3DFSC plot for the closed, **d**, 1-up and **e**, 3-up globally refined reconstructions. **f**, DeepEMhancer filtered EM density map for the closed, **g**, 1-up and **h**, 3-up *holo* HKU1-A spike ectodomains coloured according to local resolution which was calculated in cryoSPARC. **i**, Angular distribution plot calculated in cryoSPARC for particle projections in the closed, **j**, 1-up and **k**, 3-up globally refined maps.

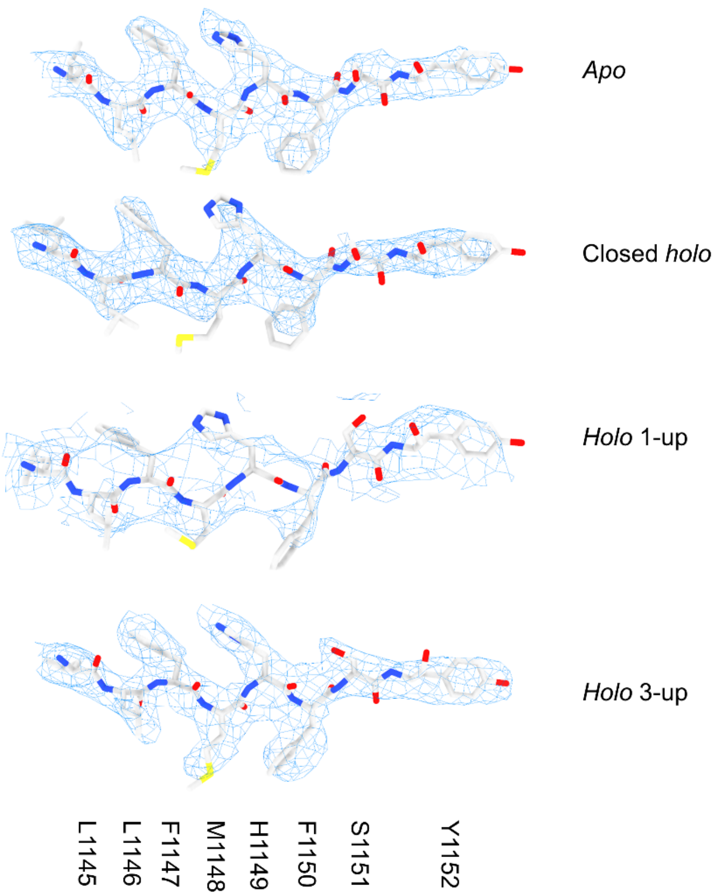

**Extended Data Fig. 12. Example density and model from the S2 region of each of the cryo-EM reconstructions generated from the *apo* and *holo* data sets.**

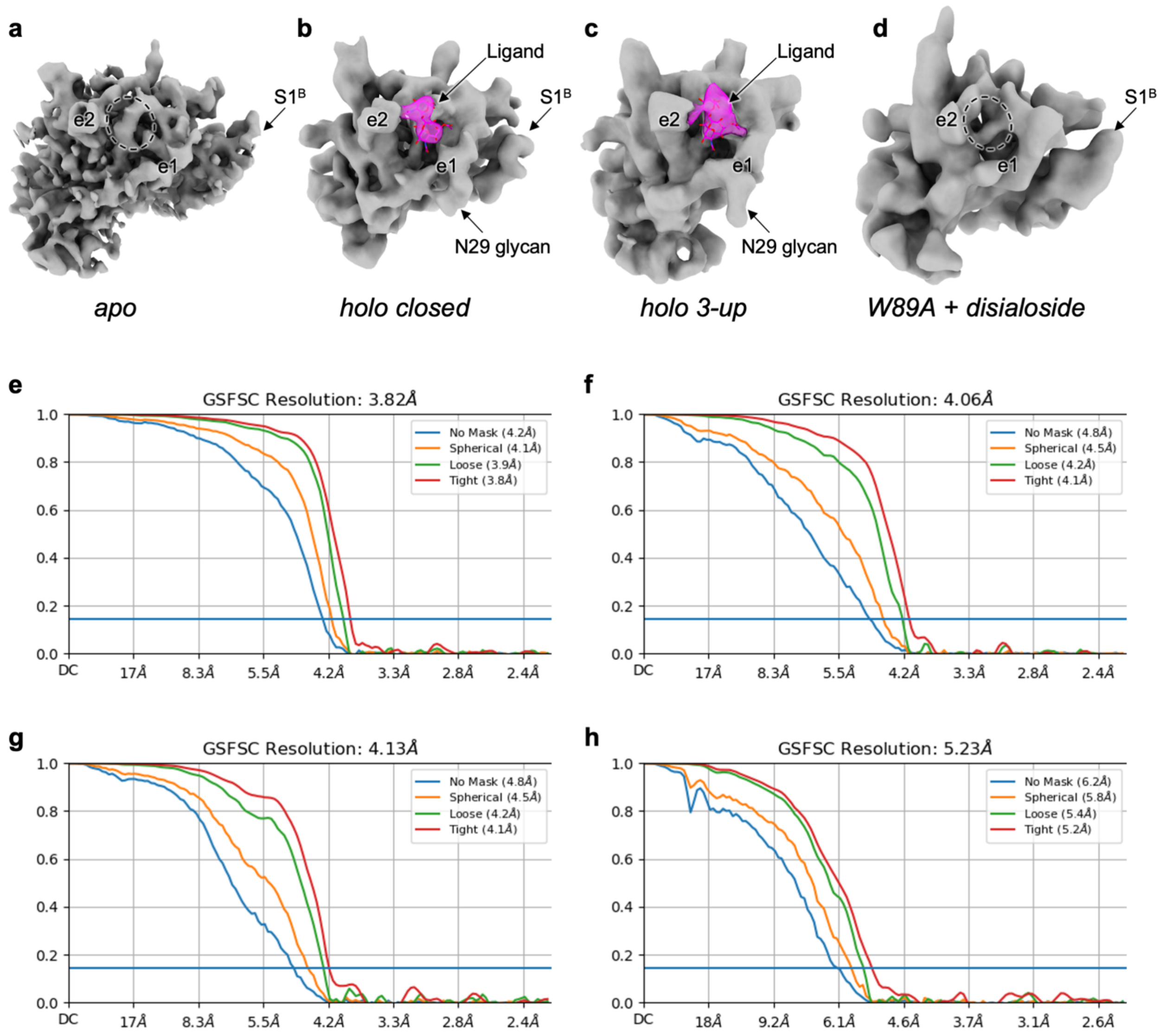

**Extended Data Fig. 13. Local refinements of the HKU1-A S1$^{A}$ domain.**

**a**, Locally refined maps of the *apo*, **b**, closed *holo*, **c**, *holo* 3-up and **d**, W89A mutant HKU1-A incubated with the disialoside. The spike protein is coloured grey and density for the disialoside, present only in the *holo* maps, is coloured magenta and contains the fitted coordinates for the molecule. In panels A and D, the receptor binding site is circled. **e**, Gold-standard FSC curves generated from the independent half maps contributing to the local refinements of the *apo*, **f**, closed *holo*, **g**, *holo* 3-up and **h**, W89A mutant HKU1-A incubated with the disialoside.

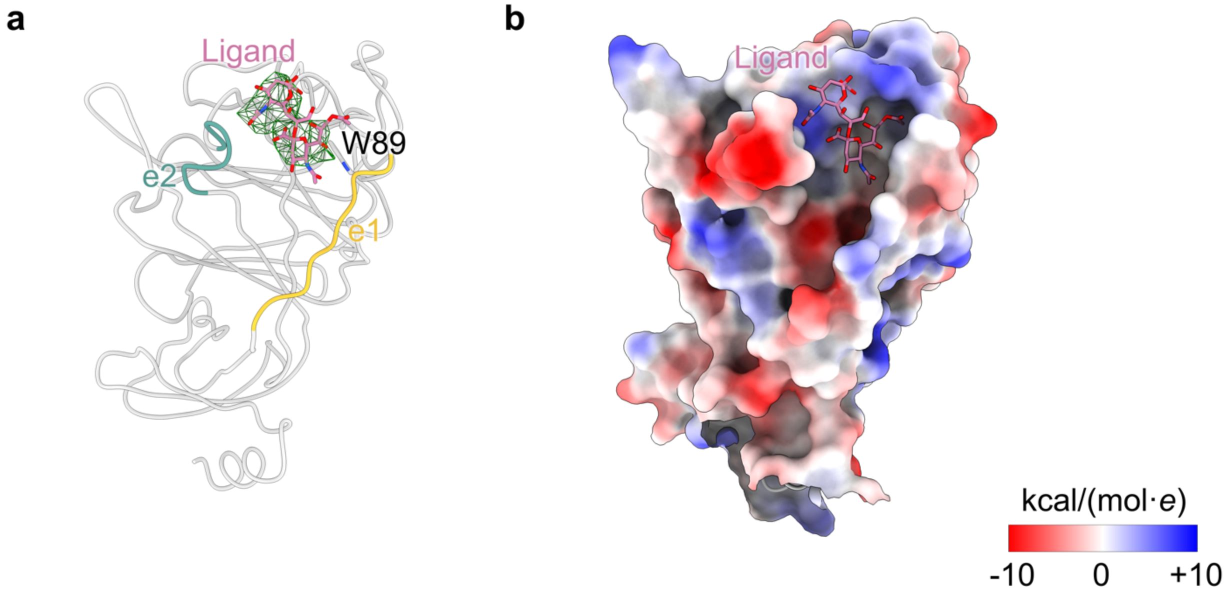

**Extended Data Fig. 14. Ligand binding site in the S1$^{A}$ domain.**

**a**, Difference density (closed *holo* minus *apo*) for the disialoside ligand bound to the S1$^{A}$ domain in the closed *holo* state confirms the binding site. **b**, Electrostatic potential map around the ligand binding site in the S1$^{A}$ domain shows a positively charged crevice in which the negatively charged disialoside binds.

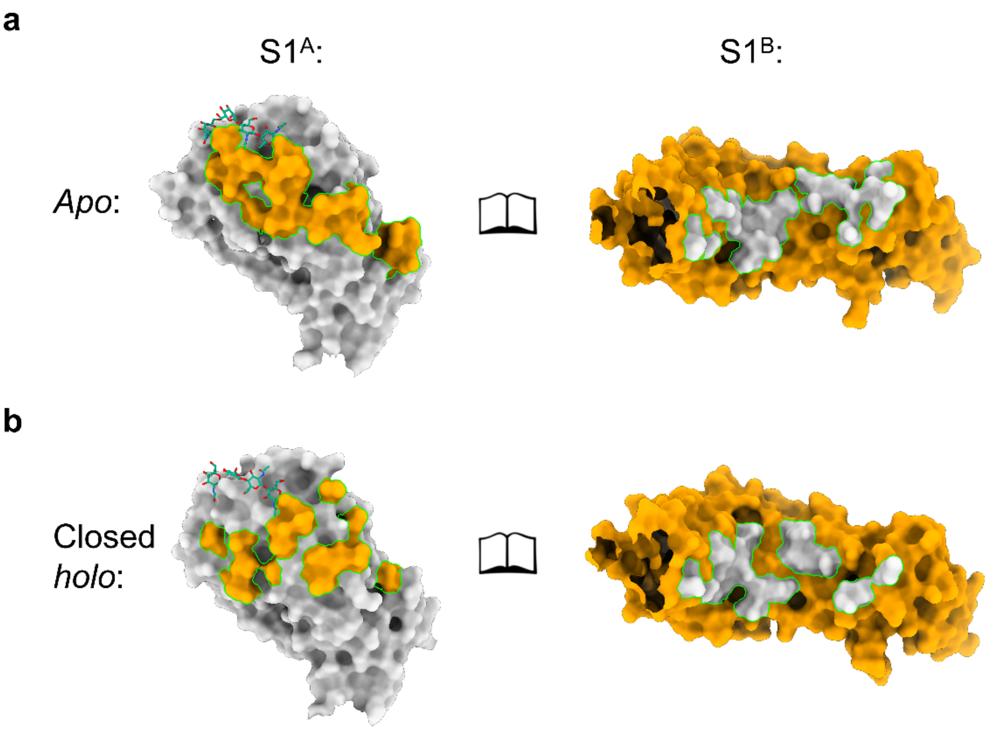

**Extended Data Fig. 15. Comparison of S1$^{A}$-S1$^{B}$ interface between *apo* and closed *holo* shows a smaller interaction footprint for the latter.**

**a**, Open book representation of the *apo* S1$^{A}$-S1$^{B}$ interface. Interacting surfaces are visualised in the colour of the subunit it interacts with. N-linked glycans on S1$^{A}$ near the interface are indicated as green sticks. **b**, *Idem* for the closed *holo* S1$^{A}$-S1$^{B}$ interface.

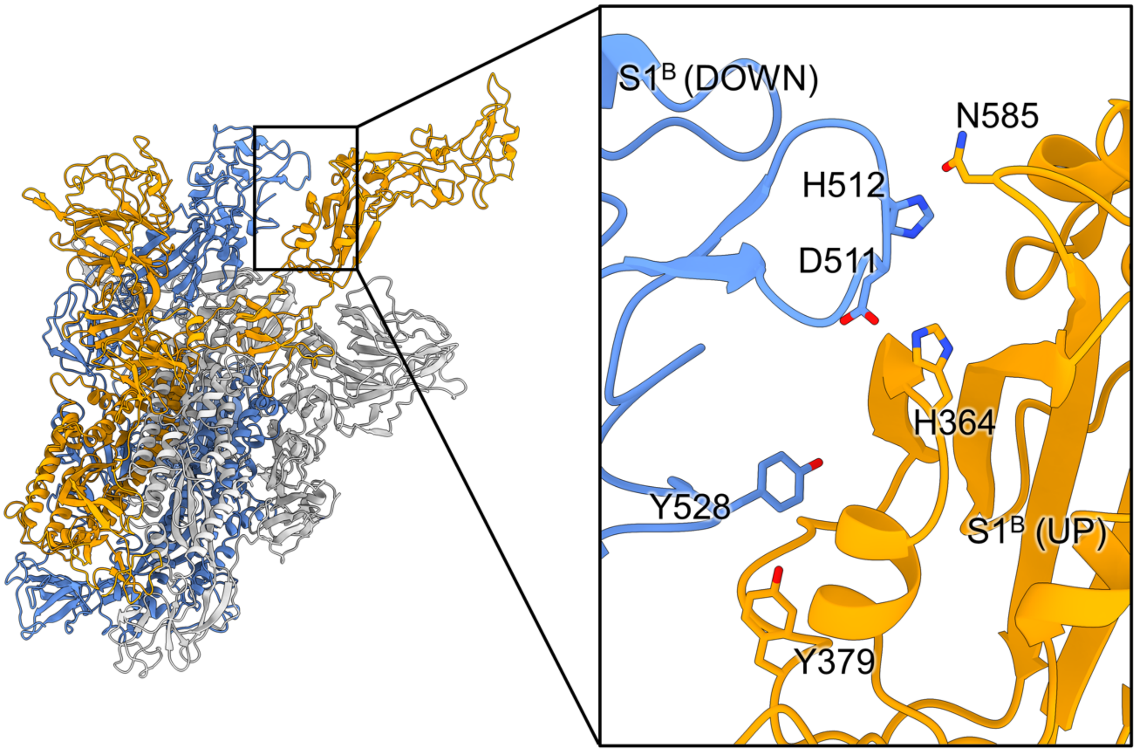

**Extended Data Fig. 16. A unique interface between the upward and downward S1$^{B}$ domains in the 1-up state.**

Residues at the interface are indicated, although the local resolution limits interpretability of side chain conformations.

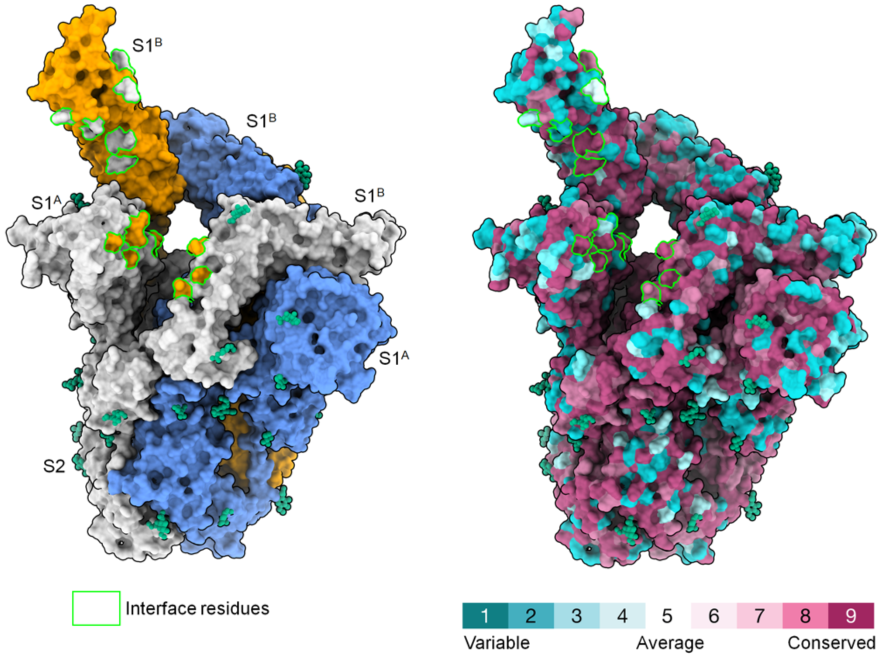

**Extended Data Fig. 17. Surface conservation of epitopes occluded in the closed state.**
In the left panel, protomers of a 1-up HKU1-A S trimer are coloured grey, blue and orange and glycans are indicated in dark green. The footprint of residues contacting neighbouring S1$^A$ and S1$^B$ domains in the closed state, but becoming exposed upon S1$^B$ flipping up, are indicated in the colour of the protomer they were originally contacting, outlined in green. The same footprints are again outlined in green in the right panel, but on the 1-up S trimer in the same orientation with its surface coloured by evolutionary conservation (glycans still indicated in dark green).

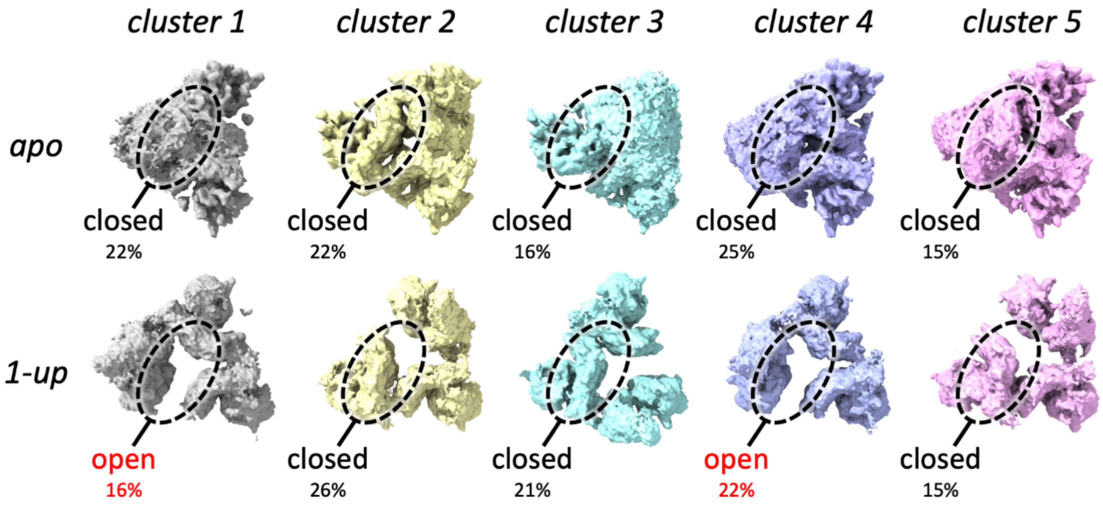

**Extended Data Fig. 18. 3D variability analysis of the *apo* and 1-up HKU1-A data sets.**
3D variability analysis of the symmetry expanded *apo* HKU1-A particles indicated that there are no detectable open S1$^B$ domains present in the data. In contrast, this method could discriminate between open and closed S1$^B$ domains in the *holo* 1-up data set, used as control to show the validity of this approach. The region which was masked during the analysis is circled.

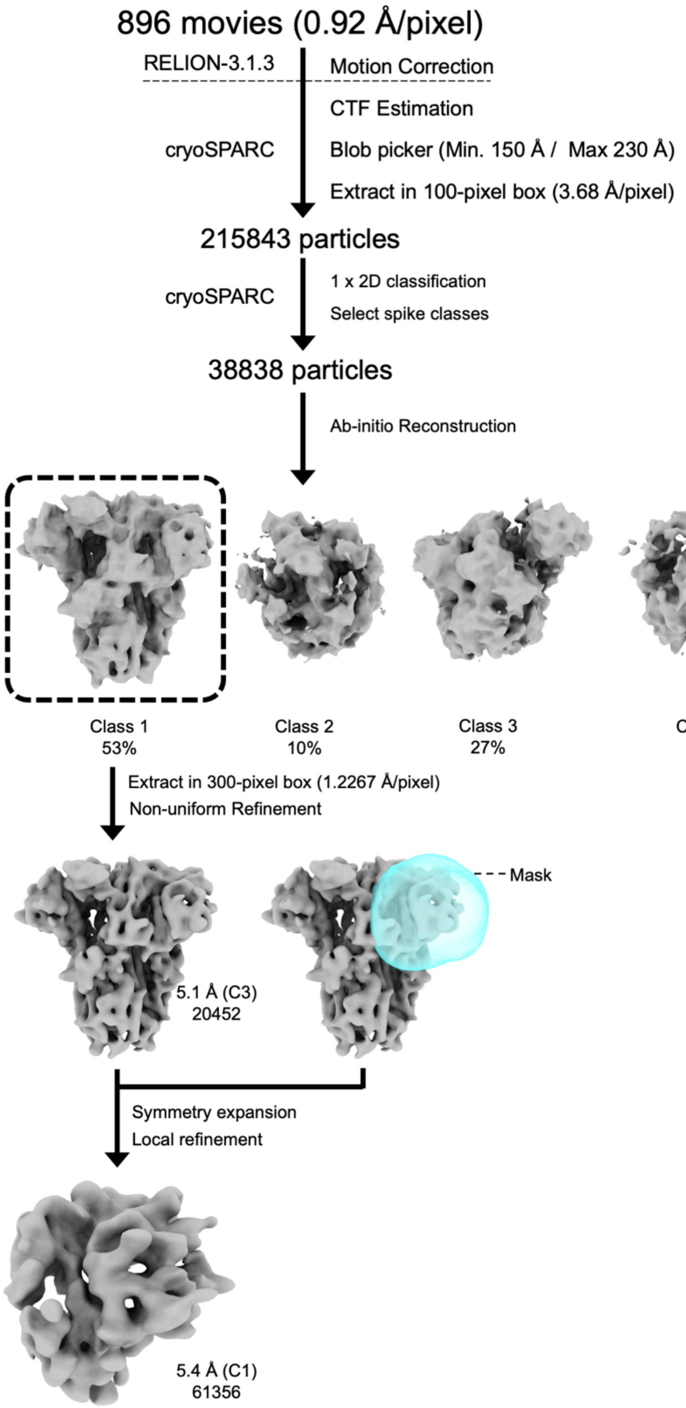

**Extended Data Fig. 19. Cryo-EM data processing pipeline for the W89A mutant HKU1-A spike glycoprotein.**

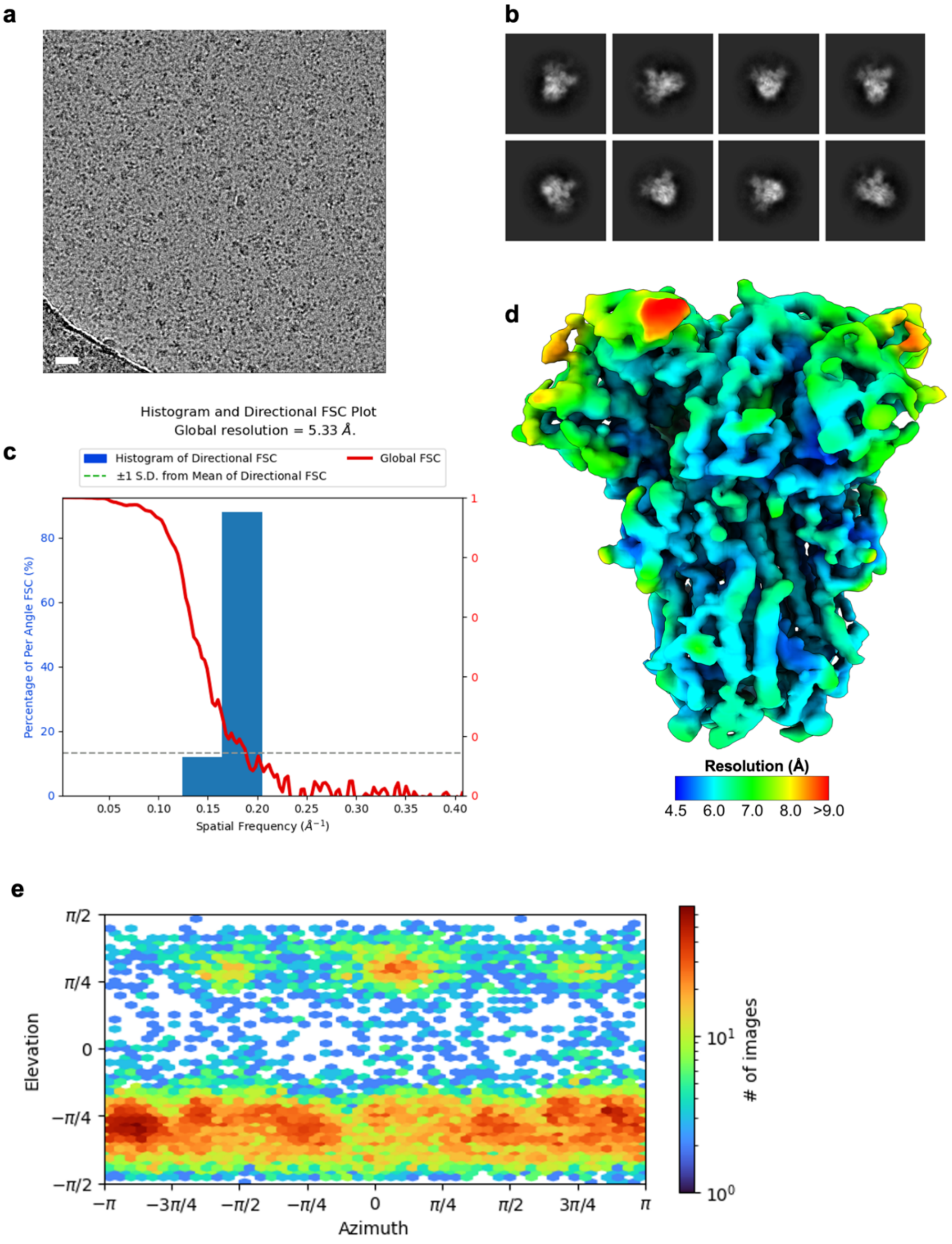

**Extended Data Fig. 20. Cryo-EM data processing of the W89A HKU1-A S ectodomain incubated with disialoside**

**a**, Representative motion-corrected micrograph of the disialoside-incubated W89A HKU1-A spike ectodomains embedded in vitreous ice. Scale bar = 50 nm. **b**, Representative reference-free 2D class averages generated in cryoSPARC. **c**, 3DFSC plot for the 5.3 Å resolution globally refined reconstruction. **d**, DeepEMhancer filtered EM density map for the *apo* HKU1-A spike ectodomains coloured according to local resolution which was calculated in cryoSPARC. **e**,

Angular distribution plot calculated in cryoSPARC for particle projections in the globally refined map.

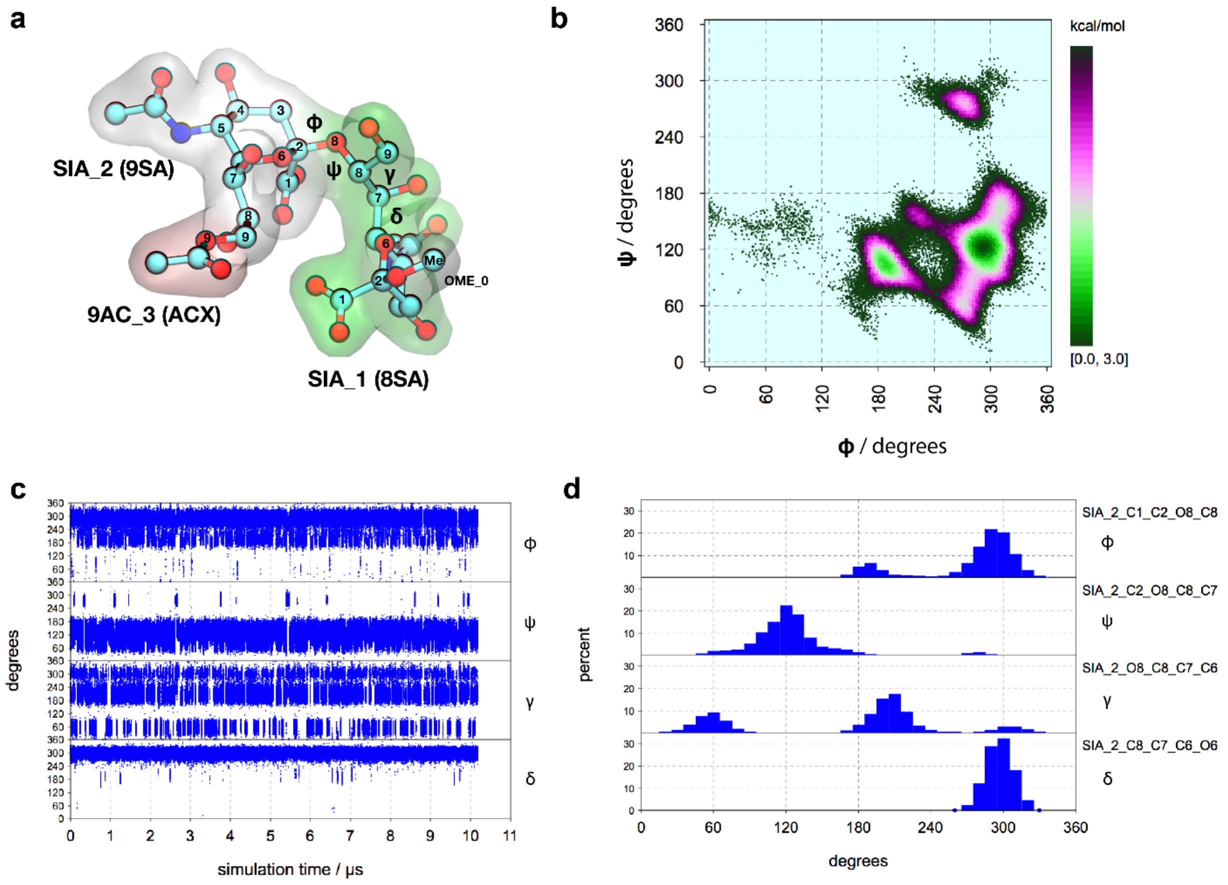

**Extended Data Fig. 21. Molecular dynamics simulations of the free disialoside.**

10 µs MD-based conformational analysis of Neu5,9Ac$_2$-α2,8-Neu5Ac-α*O*Me in explicit solvent. **a**, Example 3D structure with annotations of residue labels used (GLYCAM residue type labels are shown in brackets), atom numbering scheme and naming of the torsions describing the 2-8 linkage. The torsions are defined as φ = C1-C2-O8-C8, ψ = C2-O8-C8-C7, γ = O8-C8-C7-C6, δ = C8-C7-C6-O6. **b**, Free energy φ/ψ map. (c, d) Trajectory plots and histograms of torsions φ, ψ, γ and δ. It can be seen that conformational transitions between the population maxima (local energy minima) are fast for φ and γ. However on the 10 µs timescale only few transitions occured for ψ and δ. Torsion δ has practically only one orientation (about -60°).

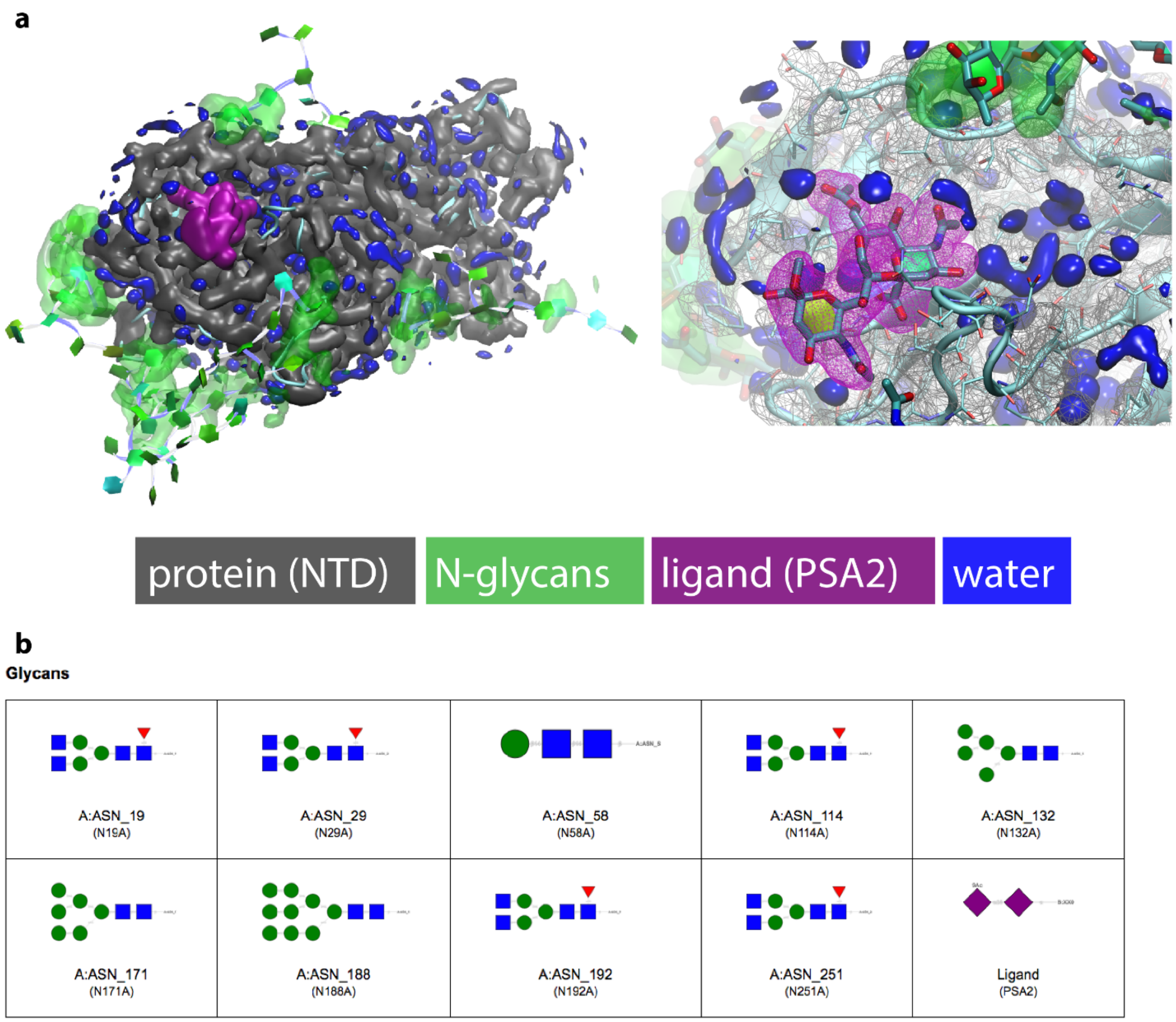

**Extended Data Fig. 22. High-resolution model of the HKU1 S (Caen1) – 2,8 disialoside complex.**

**a**, MD-derived pseudo-electron density of the disialoside ligand (PSA2, purple) in the binding pocket of the $S1^A$ domain. Data were derived from 3 µs MD simulations of $S1^A$ (NTD, residues 14-299) based on the *holo* cryo-EM model. **b**, SNFG representations of the N-glycans and the ligand. See supplementary video 6 for a 3D view of the disialoside conformation in **a**.

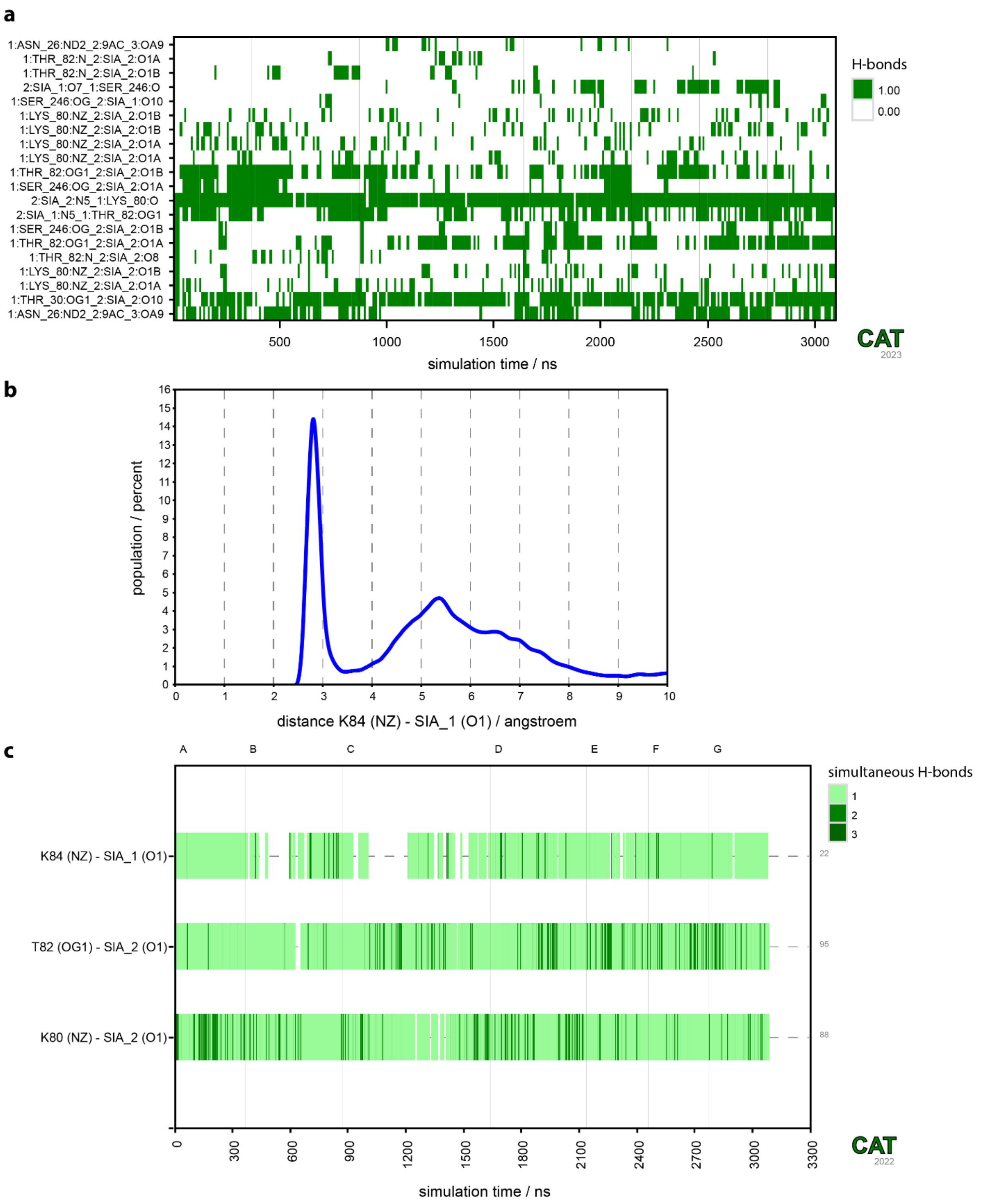

**Extended Data Fig. 23. Hydrogen bond analysis of the HKU1 S (Caen1) – α2,8 disialoside complex.**

Data were derived from 3 µs MD simulations of $S1^{A}$ (NTD, residues 14-299) based on the *holo* cryoEM model. **a**, Trajectory plots of the most populated hydrogen bonds between the ligand and the receptor protein. The existence of an H-bond at a given time is indicated in green. The H-bond labels for donor(D)-acceptor(A) atom pairs are automatically generated using a format *molD:resD:atomD_molA:resA:atomA* (see Extended Data Fig. 21 for residue names). A geometric H-bond criterion, defined as distance (D-A) ≤ 3.2 Å and angle (D-H-A) ≥ 120º, was used. **b**, Histogram of the distance K84(NZ)-SIA1(O1) showing a high probability for a salt bridge between the amino group of K84 and the carboxylate group of SIA1. **c**, Analysis of selected hydrogen bonds of the carboxyl groups of SIA1 and SIA2. Such '*complex H-bond interactions*' involve two (equivalent) acceptor atoms (O1A and O1B) and potentially multiple (equivalent) donor H-atoms (*e.g.* three in Lys:NZ), which results in multiple entries in the H-bond table (Extended Data Table 4.). The trajectory plot shows the number of individual H-bonds simultaneously formed between a donor-acceptor atom groups as a colour code. Data were analysed using Conformational Analysis Tools (CAT, http://www.md-simulations.de/CAT/).

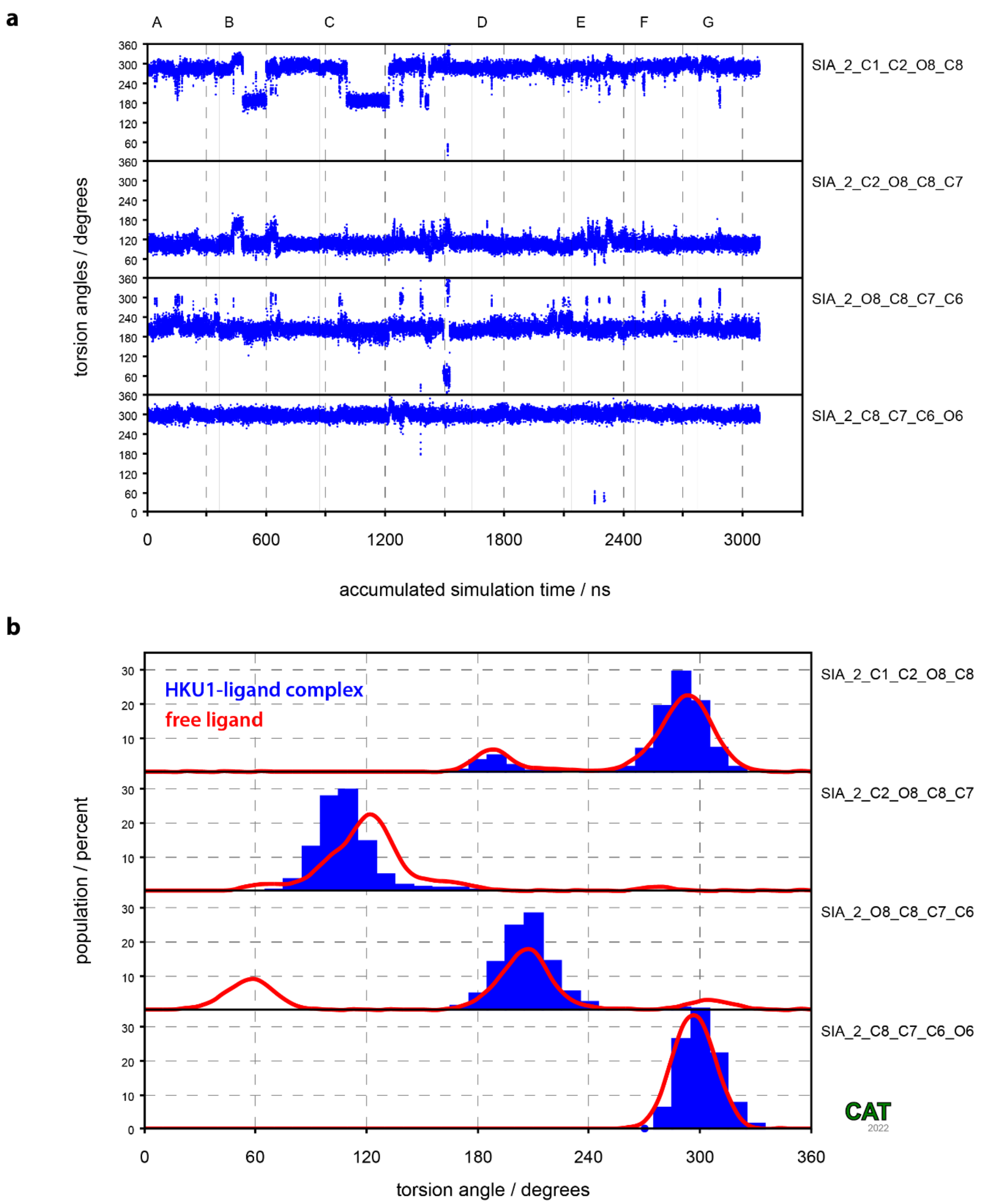

**Extended Data Fig. 24. Dynamics of the 2-8 linkage of the disialoside in the complex**

Dynamics in the 2-8 linkage are reduced but remain possible when the dialoside is bound to HKU1. **a**, Trajectory plots of linkage torsions φ, ψ, γ and δ. In comparison to the dynamics of the disaccharide in the free state, there is a clear reduction in the conformational transition frequency for torsions φ and γ (compare Extended Data Fig. 21). **b**, Histograms of linkage torsions φ, ψ, γ and δ. In comparison to the profiles of the disaccharide in the free state (red curves) there is a clear reduction in accessible conformational space for torsions γ. Whereas in the free state there are three population maxima, there is now a clear preference for a value around 210º. Data were derived from the same MD simulations shown in Extended Data Fig. 23.

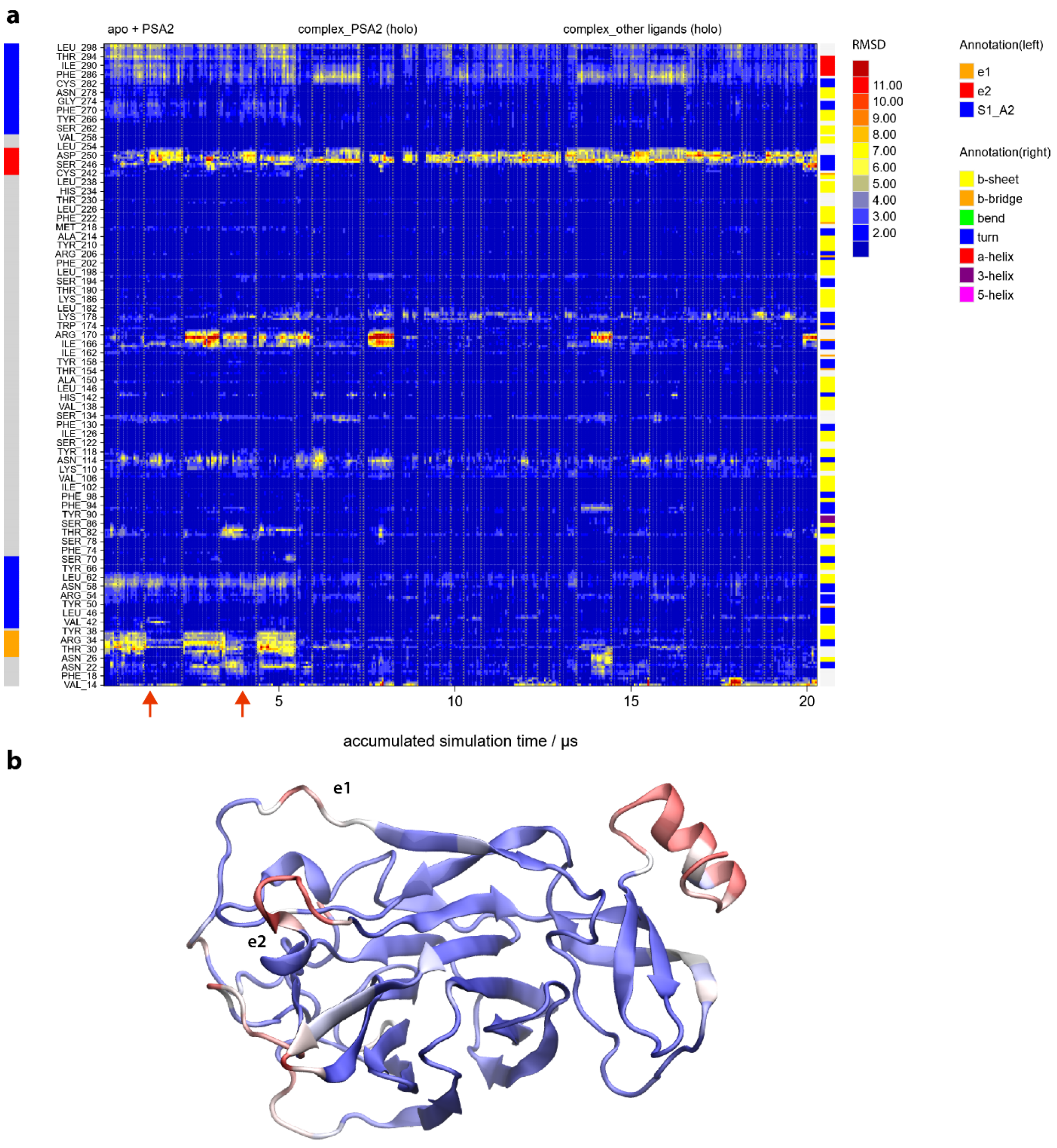

**Extended Data Fig. 25. Dynamics of HKU1 S (Caen1) $S1^A$**

In order to generate an overview of the protein dynamics over a long timescale (accumulated 20 µs), all MD simulations performed with the Caen1 sequence were combined for an RMSD analysis based on the *holo* cryoEM model as a reference structure (*i.e.* conformational changes from the *apo* into the *holo* state are apparent as transitions from high to low RMSD states). **a**, RMSD per residue trajectory plot (heat map). The individual simulations (33) are separated by vertical lines. The spontaneous conformational shifts of the e1 loop, observed in the simulations starting with the disialoside (PSA2) bound to the *apo* cryoEM conformation of the protein, are indicated with red arrows. Simulations based on the *holo* EM model with PSA2 or other Neu5,9Ac-containing ligands show ligand-induced stabilisation of the e1 conformational shift. **b**, Colour-mapping of the average RMSD values of the $C_\alpha$ atoms to the *apo* cryoEM model. Colour code: RMSD ≤ 2 Å (blue), RMSD ≥ 4 Å (red).

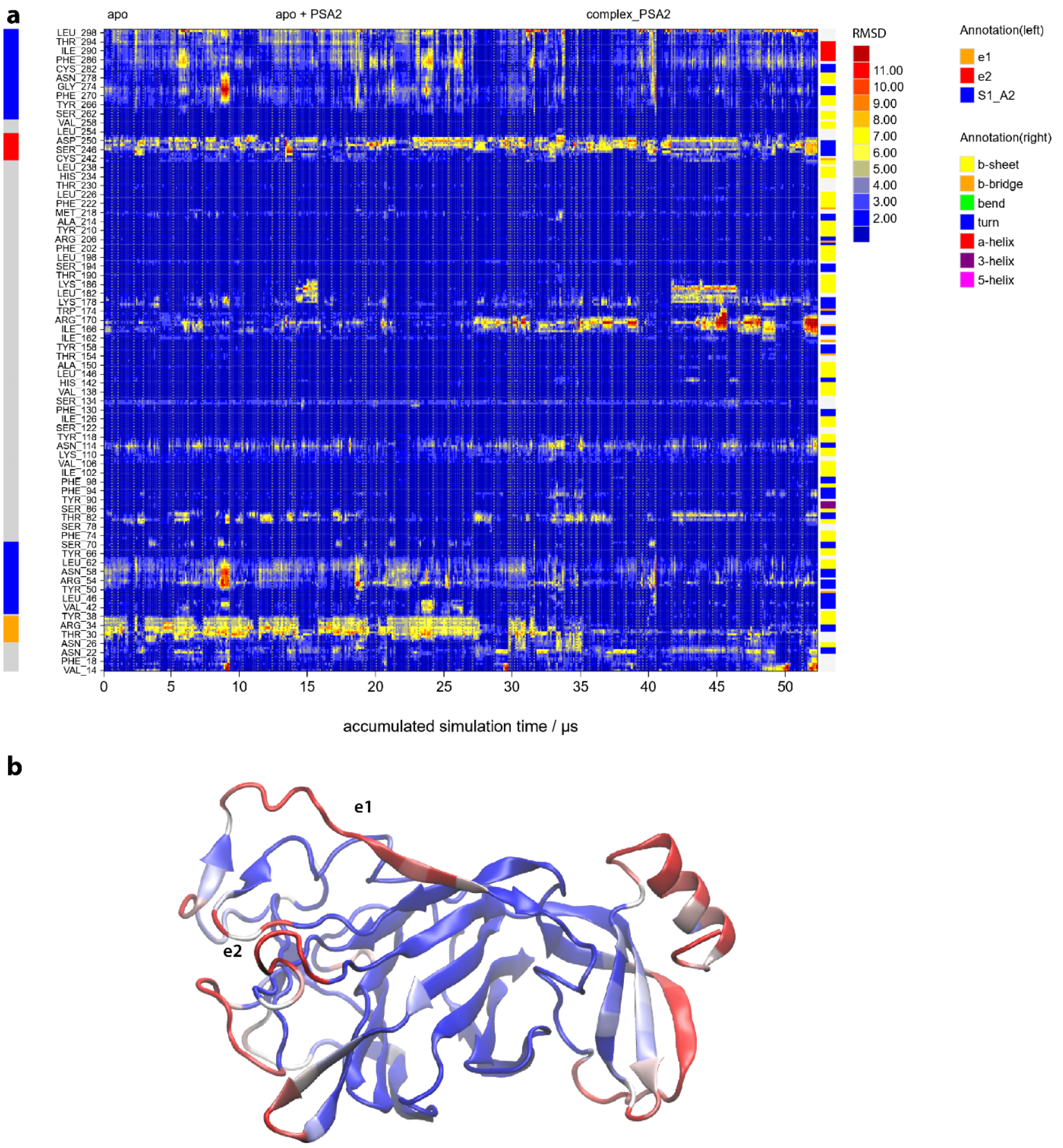

**Extended Data Fig. 26. Dynamics of HKU1 S (N1 strain) $S1^A$**

In order to generate an overview of the protein dynamics over a long timescale (accumulated 52 μs) all MD simulations performed with the N1 strain sequence were combined for an RMSD analysis based on the *holo* cryoEM model of Caen1 (replacing respective residues different in N1) as a reference structure. **a**, RMSD per residue trajectory plot (heat map). The individual simulations (69) are separated by vertical lines. **b**, Colour-mapping of the average RMSD values of the $C_\alpha$ atoms to the *apo* cryoEM model. Colour code: RMSD ≤ 2 Å (blue), RMSD ≥ 4 Å (red).

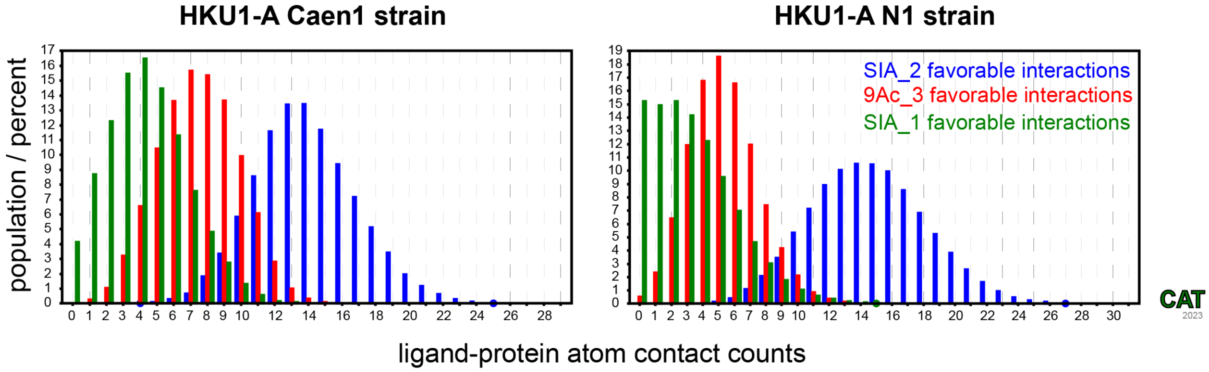

**Extended Data Fig. 27. Stabilising atom contacts between the disialoside ligand and S1$^{A}$ domains of HKU1 Caen1 or N1.**

Histogram of favourable, stabilising contacts (as defined in Extended Data Table 6) between the individual moieties of the disialoside ligand and the S1$^{A}$ domains. A notable decrease in stabilising contacts with the reducing end Sia1 can be seen in HKU1 N1, potentially due to the absence of K84 (see also the hydrogen bond analyses in Extended Data Tables 4-5).

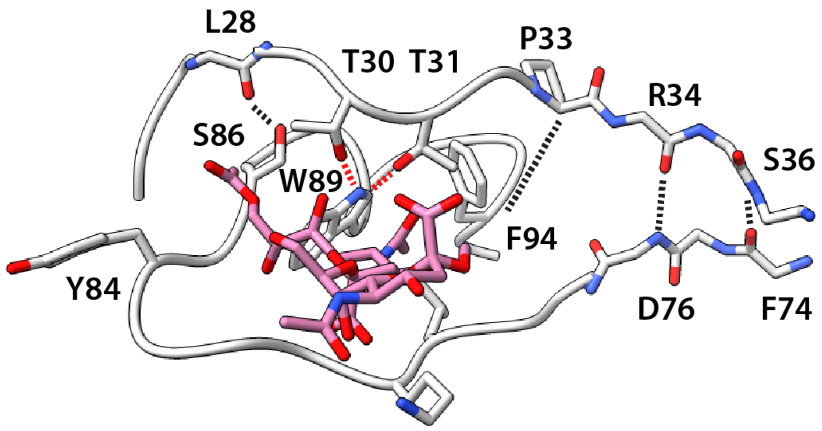

**Extended Data Fig. 28. Alternative arrangement of the disialoside binding pocket.**

The structure shown is based on MD-simulations of the HKU1-A N1 reference strain (GenBank entry NC_006577.2, Extended Data Fig. 26). Notably, the essential hydrogen bond with W89 can be formed by the T30 backbone carbonyl (*cf*. Fig. 5a), or the T30 and T31 sidechains (red lines).

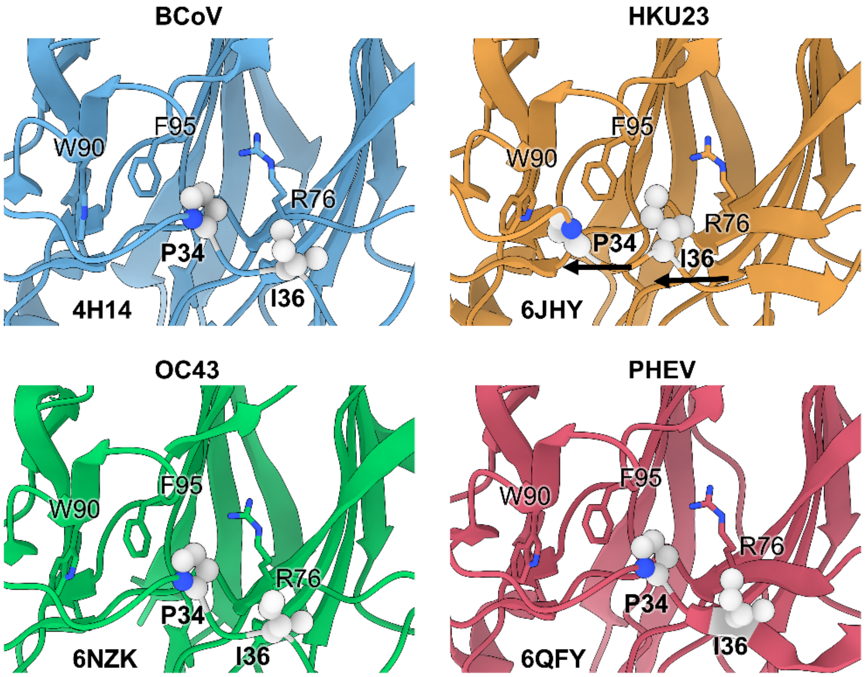

**Extended Data Fig. 29. The topology of the e1 loop in different CoVs observed in the PDB.** Shift of the e1 loop of HKU23 S[4] (orange, top-right) compared to other coronavirus S1$^{A}$ domain e1 loops[5–7], exemplified by e1 residues P34 and I36 (light grey spheres). Side chains of W90, F95 and R76 are indicated as sticks for reference. The dromedary camel CoV HKU23 is the only CoV displaying a similar conformation of e1 as found in our HKU-1 *apo* structure. We note that for BCoV, HKU23 and PHEV structures, e1 residues are involved in crystal contacts which may artificially shift the equilibrium away from preferred conformations in a physiological setting.

Amino acid sequence of HKU1 CD5-SED-GCN4-Tx-ST

MPMGSLQPLATLYLLGMLVASVLAVIGDFNCTNFAINDLNTTIPRISEYVVDVSYGLGTYYILDRVYLNT
TILFTGYFPKSGANFRDLSLKGTTKLSTLWYQKPFLSDFNNGIFSRVKNTKLYVNKTLYSEFSTIVIGSV
FINNSYTIVVQPHNGVLEITACQYTMCEYPHTICKSIGSSRNESWHFDKSEPLCLFKKNFTYNVSTDWLY
FHFYQERGTFYAYYADSGMPTTFLFSLYLGTLLSHYYVLPLTCNAISSNTDNETLQYWVTPLSKRQYLLK
FDDRGVITNAVDCSSSFFSEIQCKTKSLLPNTGVYDLSGFTVKPVATVHRRIPDLPDCDIDKWLNNFNVP
SPLNWERKIFSNCNFNLSTLLRLVHTDSFSCNNFDESKIYGSCFKSIVLDKFAIPNSRRSDLQLGSSGFL
QSSNYKIDTTSSSCQLYYSLPAINVTINNYNPSSWNRRYGFNNFNLSSHSVVYSRYCFSVNNTFCPCAKP
SFASSCKSHKPPSASCPIGTNYRSCESTTVLDHTDWCRCSCLPDPITAYDPRSCSQKKSLVGVGEHCAGF
GVDEEKCGVLDGSYNVSCLCSTDAFLGWSYDTCVSNNRCNIFSNFILNGINSGTTCSNDLLQPNTEVFTD
VCVDYDLYGITGQGIFKEVSAVYYNSWQNLLYDFNGNIIGFKDFVTNKTYNIFPCYAGRVSAAFHQNASS
LALLYRNLKCSYVLNNISLATQPYFDSYLGCVFNADNLTDYSVSSCALRMGSGFCVDYNSPSSSSSGGSG
SSISASYRFVTFEPFNVSFVNDSIESVGGLYEIKIPTNFTIVGQEEFIQTNSPKVTIDCSLFVCSNYAAC
HDLLSEYGTFCDNINSILDEVNGLLDTTQLHVADTLMQGVTLSSNLNTNLHFDVDNINFKSLVGCLGPHC
GSSSRSFFEDLLFDKVKLSDVGFVEAYNNCTGGSEIRDLLCVQSFNGIKVLPPILSESQISGYTTAATVA
AMFPPWSAAAGIPFSLNVQYRINGLGVTMDVLNKNQKLIATAFNNALLSIQNGFSATNSALAKIQSVVNS
NAQALNSLLQQLFNKFGAISSSLQEILSRLDALEAQVQIDRLINGRLTALNAYVSQQLSDISLVKLGAAL
AMEKVNECVKSQSPRINFCGNGNHILSLVQNAPYGLLFMHFSYKPISFKTVLVSPGLCISGDVGIAPKQG
YFIKHNDHWMFTGSSYYYPEPISDKNVVFMNTCSVNFTKAPLVYLNHSVPKLSDFESELSHWFKNQTSIA
PNLTLNLHTINATFLDLLIKRMKQIEDKIEEIESKQKKIENEIARIKKIKLVPRGSLEWSHPQFEK*

Coding sequence HKU1 CD5-SED-GCN4-Tx-ST

**ATGCCCATGGGGTCTCTGCAACCGCTGGCCACCTTGTACCTGCTGGGGATGCTGGTCGCTTCCGTGCTA**g
caGTTATAGGTGATTTTAATTGTACTAATTTTGCTATTAATGATTTAAACACCACAATTCCTCGCATAAG
TGAGTATGTTGTGGATGTTTCTTATGGTTTGGGTACATATTATATACTTGATCGTGTTTATTTAAATACT
ACTATATTATTTACTGGTTATTTCCCTAAATCTGGTGCCAATTTTAGGGATCTATCTTTAAAAGGTACTA
CAAAATTGAGTACTCTTTGGTATCAGAAACCCTTTTTATCTGATTTTAATAATGGTATTTTTTCTAGAGT
TAAGAATACTAAGTTGTATGTTAATAAAACTTTGTATAGTGAGTTTAGTACTATAGTTATAGGTAGTGTT
TTTATTAACAACTCTTATACTATTGTTGTTCAACCTCATAATGGTGTTTTGGAGATTACAGCTTGTCAAT
ACACTATGTGTGAGTATCCTCATACTATTTGTAAATCTATAGGTAGTTCTCGTAATGAATCTTGGCATTT
TGATAAATCTGAACCTTTGTGTCTGTTCAAGAAAAATTTTACTTATAATGTTTCTACAGATTGGTTGTAT
TTTCATTTTTATCAAGAACGTGGCACTTTTTATGCTTATTATGCTGATTCTGGCATGCCTACTACTTTTT
TATTTAGTTTGTATCTTGGTACTCTTTTATCTCATTATTATGTTTTGCCTTTGACTTGTAATGCTATATC
TTCTAATACTGATAATGAGACTTTACAATATTGGGTCACACCTTTGTCTAAACGCCAATATCTTCTTAAA
TTTGACGACCGTGGTGTTATTACTAATGCTGTTGATTGTTCTAGTAGTTTCTTTAGCGAGATTCAATGTA
AAACTAAATCTTTATTACCTAATACTGGTGTTTATGACTTATCTGGTTTTACTGTTAAGCCTGTTGCAAC
TGTACATCGTCGTATTCCTGATTTACCTGATTGTGACATTGATAAATGGCTTAACAATTTTAATGTACCC
TCACCTCTTAATTGGGAACGTAAAATTTTTTCTAATTGCAACTTTAATTTGAGTACTTTGCTTCGTTTAG
TTCATACTGATTCTTTTTCTTGTAATAATTTTGATGAATCTAAGATATATGGTAGTTGTTTTAAGAGTAT
TGTTTTAGATAAATTTGCCATACCCAACTCCAGACGATCTGATTTGCAGTTGGGCAGTTCTGGTTTTCTG
CAATCTTCTAATTATAAAATTGACACTACTTCTAGTTCTTGTCAATTGTATTATAGTTTGCCTGCAATTA
ATGTTACTATTAATAATTATAATCCTTCTTCTTGGAATAGAAGGTATGGTTTTAATAATTTTAATTTGAG
TTCTCATAGTGTTGTTTACTCACGTTATTGTTTTTCTGTTAATAATACTTTTTGTCCTTGTGCTAAACCT
TCTTTTGCTTCAAGTTGCAAGAGTCATAAACCACCTTCTGCTTCTTGTCCTATTGGTACTAATTATCGTT
CTTGTGAGAGTACTACTGTACTCGACCACACTGACTGGTGTAGGTGTTCTTGTTTACCTGATCCTATAAC
TGCTTATGACCCTAGGTCTTGTTCTCAAAAAAAGTCTCTGGTTGGTGTTGGTGAACATTGTGCAGGGTTC
GGTGTTGATGAAGAAAAGTGTGGTGTATTGGATGGATCATATAATGTTTCTTGTCTTTGTAGTACTGATG
CCTTTCTAGGTTGGTCTTATGACACTTGCGTCAGTAACAACCGTTGTAATATTTTTTCTAATTTTATTTT
AAATGGTATCAATAGTGGTACCACTTGTTCTAATGATTTATTGCAGCCTAATACTGAAGTTTTTACTGAT
GTTTGTGTTGATTACGACCTTTATGGTATTACAGGACAAGGTATTTTTAAAGAAGTTTCTGCTGTTTATT

ATAATAGTTGGCAAAATCTTTTGTATGATTTTAATGGCAACATTATTGGTTTTAAAGATTTTGTTACTAA
TAAAACATATAATATTTTCCCTTGTTATGCAGGAAGAGTTTCTGCTGCTTTTCATCAAAATGCTTCCTCT
TTGGCTTTACTTTATCGTAATTTAAAATGTAGCTATGTTTTGAATAATATTTCTTTAGCTACTCAGCCAT
ATTTTGATAGTTATCTTGGTTGCGTTTTTAATGCTGATAATTTAACTGATTATTCTGTTTCTTCTTGTGC
TCTTCGCATGGGTAGTGGTTTTTGTGTTGATTATAACTCACCTTCTTCTTCCTCTTCGGGTGGTTCTGGT
TCTAGTATTTCTGCTTCTTATCGGTTTGTTACTTTTGAACCCTTTAATGTCAGTTTTGTTAATGACAGTA
TTGAGTCTGTGGGTGGTCTTTATGAGATCAAAATTCCCACTAACTTTACTATAGTTGGTCAAGAGGAATT
TATTCAAACTAATTCTCCTAAAGTTACTATTGATTGTTCTTTATTTGTCTGTTCTAATTATGCAGCTTGC
CATGACTTATTGTCAGAGTATGGCACTTTTTGTGATAATATTAATAGTATTTTAGATGAAGTTAATGGTT
TACTTGATACTACTCAATTGCATGTAGCTGATACTCTTATGCAAGGTGTCACACTTAGCTCCAATCTTAA
TACTAATTTGCATTTTGATGTTGATAATATTAATTTTAAATCCCTAGTTGGATGTTTAGGTCCACACTGC
GGTTCTTCTTCTCGTTCTTTTTTTGAAGATTTATTGTTTGACAAAGTTAAACTTTCAGATGTTGGTTTTG
TTGAAGCTTATAACAATTGTACTGGTGGTAGTGAAATTAGAGATCTTCTTTGTGTACAATCCTTTAATGG
TATTAAAGTTTTGCCTCCTATTTTGTCTGAATCTCAAATTTCTGGTTACACCACAGCCGCTACTGTTGCT
GCTATGTTTCCACCATGGTCAGCAGCAGCTGGCATACCATTTTCTCTTAATGTACAATATAGAATTAATG
GTTTGGGTGTTACTATGGATGTTCTTAATAAAAATCAAAAGTTGATAGCTACTGCTTTTAATAATGCTCT
TCTTTCTATTCAGAATGGTTTTAGTGCTACCAACTCTGCACTTGCTAAAATACAAAGTGTTGTTAATTCT
AATGCTCAAGCACTTAATAGTTTGTTACAGCAATTATTTAATAAATTTGGTGCAATTAGTTCTTCTTTAC
AAGAAATTTTATCTCGTCTCGATGCTTTAGAGGCTCAGGTTCAGATTGATAGGCTTATTAATGGTCGTTT
AACTGCTTTAAATGCTTATGTTTCTCAACAGCTTAGTGATATTTCTCTTGTAAAACTTGGTGCTGCTTTA
GCTATGGAGAAGGTTAATGAGTGTGTTAAAAGTCAATCTCCTCGTATTAATTTTTGTGGTAATGGTAATC
ATATTTTGTCATTAGTTCAAAATGCTCCTTATGGTTTGTTGTTTATGCATTTTAGTTATAAACCTATTTC
TTTTAAAACTGTTTTAGTAAGTCCTGGTTTATGTATATCAGGTGATGTAGGTATTGCACCTAAACAAGGG
TATTTTATTAAACATAATGATCATTGGATGTTTACTGGTAGTTCTTACTATTATCCTGAACCAATTTCAG
ATAAAAATGTTGTTTTTATGAATACTTGTTCTGTTAATTTTACTAAAGCGCCTCTTGTTTATTTGAATCA
TTCTGTACCAAAATTGTCTGATTTTGAATCTGAGTTATCTCATTGGTTTAAAAATCAAACATCCATTGCG
CCTAATTTGACTTTAAATCTTCATACTATTAATGCTACTTTTTTAGATTTGttaATTAAgCGCATGAAGC
AGATCGAGGACAAGATCGAAGAGATCGAGTCCAAGCAGAAGAAGATCGAGAACGAGATCGCCCGCATCAA
GAAGattaagctggtgccgcgcggcagcctcgagtggagccacccgcagttcgagaagtga

**Extended Data Fig. 30. HKU1-A S-ectodomain construct used in this study.**

The signal peptide (yellow), GCN4 trimerization domain (green), thrombin cleavage site (purple) and Strep-Tag (red) are indicated.

**Extended Data Table 1. Summary of coronavirus spike proteins determined by cryo-EM and their $S1^B$ state.**

| Subgenus | Virus | PDB ID | Reference | $S1^B$ up? |
|---|---|---|---|---|
| **Alpha** | | | | |
| *Duvinacovirus* | HCoV-229E | *6U7H, 7CYC* | 10.7554/eLife.51230[8]<br>10.1038/s41467-020-20401-y[9] | No |
| *Setracovirus* | HCoV-NL63 | *5SZS* | 10.1038/nsmb.3293[10] | No |
| *Tegacovirus* | FIPV | *6JX7* | 10.1073/pnas.1908898117[11] | No |
| | CCoV-HuPn-2018 | *7U0L, 7US6, 7US9, 7USA, 7USB* | 10.1016/j.cell.2022.05.019[12] | No |
| *Rhinacovirus* | HKU2 | *6M15* | 10.1038/s41467-020-16876-4[13] | No |
| | SADS-CoV | *6M39* | 10.1128/JVI.01301-20[14] | No |
| *Pedacovirus* | PEDV | *6U7K, 6VV5, 7W6M, 7W73, 7Y6S, 7Y6T, 7Y6U, 7Y6V* | 10.1038/s41467-022-32588-3[15]<br>10.1016/j.str.2020.12.003[16]<br>10.1128/JVI.00923-19[17] | *Yes* |
| **Beta** | | | | |
| *Embecovirus* | MHV | *3JCL, 6VSJ* | 10.1038/nature16988[18]<br>10.1371/journal.ppat.1008392[19] | No |
| | HCoV-HKU1-B | *5I08* | 10.1038/nature17200[3] | No |
| | HCoV-OC43 | *6OHW, 6NZK, 7SB3* | 10.1038/s41594-019-0233-y[6]<br>10.1126/sciadv.abn2911[20] | No |
| *Sarbecovirus* | SARS-CoV | *5X5B, 5X58* | 10.1038/ncomms15092[21] | *Yes* |
| | SARS-CoV-2 | *6VXX, 6VYB* | 10.1016/j.cell.2020.02.058[22] | *Yes* |
| | Pangolin sarbecovirus | *7BBH, 7CN8* | 10.1038/s41467-021-21006-9[23]<br>10.1038/s41467-021-21767-3[24] | No |
| | RaTG13 | *6ZGF* | 10.1038/s41594-020-0468-7[25] | No |
| *Merbecovirus* | MERS-CoV | *5X59, 5X5F* | 10.1038/ncomms15092[21] | *Yes* |
| | PDF-2180 | *7U6R* | 10.1038/s41586-022-05513-3[26] | No |
| **Gamma** | | | | |
| *Igacovirus* | IBV | *6CV0* | 10.1371/journal.ppat.1007009[27] | No |
| **Delta** | | | | |
| *Buldecovirus* | PDCoV | *6BFU, 6B7N* | 10.1128/JVI.01628-17[28]<br>doi.org/10.1128/JVI.01556-17[29] | No |

**Extended Data Table 2. Cryo-EM data collection, refinement and validation statistics for global refinements.**

| | *apo* (EMDB-16882) (PDB 8OHN) | closed *holo* (EMDB-17076) (PDB 8OPM) | *holo* 1-up (EMDB-17077) (PDB 8OPN) | *holo* 3-up (EMDB-17078) (PDB 8OPO) | *holo* mutant W89A (EMDB-17079) |
|---|---|---|---|---|---|
| **Data collection and processing** | | | | | |
| Magnification | 105,000x | 105,000x | 105,000x | 105,000x | 150,000x |
| Voltage (kV) | 300 | 300 | 300 | 300 | 200 |
| Electron exposure (e–/Å$^2$) | 46.3 | 46.3 | 46.3 | 46.3 | 41.7 |
| Defocus range (μm) | 1.5-2.5 | 1.5-2.5 | 1.5-2.5 | 1.5-2.5 | 1.5-2.5 |
| Pixel size (Å) | 0.415* | 0.415* | 0.415* | 0.415* | 0.92 |
| Symmetry imposed | C3 | C3 | C1 | C3 | C3 |
| Initial particle images (no.) | 914772 | 956697 | 956697 | 956697 | 215843 |
| Final particle images (no.) | 108396 | 44081 | 36048 | 99174 | 38838 |
| Map resolution (Å) | 3.4 | 3.8 | 5 | 3.7 | 5.3 |
| FSC threshold | 0.143 | 0.143 | 0.143 | 0.143 | 0.143 |
| Map resolution range (Å) | 2.4-11 | 3.2-13 | 4.1-17 | 2.4-12 | 4.6-12.7 |
| **Refinement** | | | | | |
| Initial model used (PDB code) | 5KWB, 6NZK | 5KWB, 6NZK | 5KWB, 6NZK | 5KWB, 6NZK | - |
| Model resolution (Å) FSC threshold 0.5 | 3.7 | 4.1 | 6.0 | 4.1 | - |
| Map sharpening *B* factor (Å$^2$) | 105 | 165 | 300 | 55 | - |
| Model composition | | | | | |
| Non-hydrogen atoms | 29373 | 29802 | 28543 | 28965 | - |
| Protein residues | 3585 | 3582 | 3555 | 3585 | - |
| Ligands | 90 | 123 | 40 | 57 | - |
| *B* factors (Å$^2$) | | | | | |
| Protein | 113.75 | 171.11 | 368.13 | 28.87 | - |
| Ligand | 144.64 | 226.06 | 396.36 | 63.10 | - |
| R.m.s. deviations | | | | | |
| Bond lengths (Å) | 0.003 | 0.004 | 0.004 | 0.003 | - |
| Bond angles (°) | 0.674 | 0.816 | 0.907 | 0.820 | - |
| Validation | | | | | |
| MolProbity score | 1.50 | 1.84 | 1.86 | 1.67 | - |
| Clashscore | 4.10 | 6.42 | 7.68 | 5.52 | - |
| Poor rotamers (%) | 0.00 | 0.00 | 0.00 | 0.00 | - |
| Ramachandran plot | | | | | |
| Favored (%) | 95.55 | 91.90 | 93.27 | 94.65 | - |
| Allowed (%) | 4.45 | 7.93 | 6.59 | 5.26 | - |
| Disallowed (%) | 0.00 | 0.17 | 0.14 | 0.08 | - |

*Super-resolution pixel size

**Extended Data Table 3. Cryo-EM data collection and refinement statistics for local refinements.**

| | *apo* (EMDB-17080) | closed *holo* (EMDB-17081) | *holo* 3-up (EMDB-17082) | *holo* mutant W89A (EMDB-17083) |
|---|---|---|---|---|
| **Data collection and processing** | | | | |
| Magnification | 105,000x | 105,000x | 105,000x | 150,000x |
| Voltage (kV) | 300 | 300 | 300 | 200 |
| Electron exposure (e–/Å$^2$) | 46.3 | 46.3 | 46.3 | 41.7 |
| Defocus range (μm) | 1.5-2.5 | 1.5-2.5 | 1.5-2.5 | 1.5-2.5 |
| Pixel size (Å) | 0.415* | 0.415* | 0.415* | 0.92 |
| Symmetry imposed | C1 | C1 | C1 | C1 |
| Initial particle images (no.) | 914772 | 956697 | 956697 | 215843 |
| Final particle images (no.) | 108396 | 71458 | 99174 | 61356 |
| Map resolution (Å) | 3.8 | 4.1 | 4.1 | 5.2 |
| FSC threshold | 0.143 | 0.143 | 0.143 | 0.143 |
| Map resolution range (Å) | 3.2-12.5 | 3.5-9 | 3.5-9.2 | 4.6-16.4 |

*Super-resolution pixel size

**Extended Data Table 4. HKU1 Caen1 $S1^A$-ligand hydrogen bond analysis.**
H-bonds were identified based on the criteria given in Extended Data Fig. 23. Distances were calculated between heavy atoms, angles were measured between donor, H, and acceptor.

| Index | Donor | Acceptor | Population | Distance | Angle |
|---|---|---|---|---|---|
| 1 | B:SIA_2:N5(HN5) | A:LYS_80:O | 96.9 | 2.86 | 156.9 |
| 2 | A:THR_30:OG1(HG1) | B:SIA_2:O10 | 80.3 | 2.82 | 159.9 |
| 3 | A:THR_82:OG1(HG1) | B:SIA_2:O1B | 58.0 | 2.75 | 157.5 |
| 4 | B:SIA_1:N5(HN5) | A:THR_82:OG1 | 56.0 | 2.99 | 157.4 |
| 5 | A:THR_82:OG1(HG1) | B:SIA_2:O1A | 49.8 | 2.76 | 158.5 |
| 6 | A:ASN_26:ND2(HD21) | B:9AC_3:OA9 | 46.8 | 2.97 | 158.9 |
| 7 | A:SER_246:OG(HG) | B:SIA_2:O1A | 23.0 | 2.68 | 160.5 |
| 8 | B:SIA_1:O7(HO7) | A:SER_246:O | 21.1 | 2.77 | 161.0 |
| 9 | A:LYS_80:NZ(HZ3) | B:SIA_2:O1A | 18.3 | 2.85 | 151.7 |
| 10 | A:LYS_80:NZ(HZ2) | B:SIA_2:O1A | 18.1 | 2.84 | 151.2 |
| 11 | A:LYS_80:NZ(HZ3) | B:SIA_2:O1B | 18.0 | 2.86 | 148.5 |
| 12 | A:LYS_80:NZ(HZ2) | B:SIA_2:O1B | 16.9 | 2.84 | 149.4 |
| 13 | A:LYS_80:NZ(HZ1) | B:SIA_2:O1B | 16.5 | 2.84 | 150.3 |
| 14 | A:LYS_80:NZ(HZ1) | B:SIA_2:O1A | 15.4 | 2.84 | 152.1 |
| 15 | A:SER_246:OG(HG) | B:SIA_2:O1B | 14.5 | 2.68 | 159.9 |
| 16 | A:THR_82:N(H) | B:SIA_2:O1B | 9.5 | 2.90 | 144.6 |
| 17 | A:ASN_26:ND2(HD22) | B:9AC_3:OA9 | 6.9 | 2.91 | 157.4 |
| 18 | A:THR_82:N(H) | B:SIA_2:O8 | 6.9 | 3.03 | 154.8 |
| 19 | A:THR_82:N(H) | B:SIA_2:O1A | 6.5 | 2.90 | 146.0 |
| 20 | A:SER_246:OG(HG) | B:SIA_1:O10 | 5.2 | 2.76 | 158.8 |
| 21 | B:SIA_1:O7(HO7) | A:SER_246:OG | 4.6 | 2.91 | 151.7 |
| 22 | A:LYS_84:NZ(HZ1) | B:SIA_1:O1A | 4.5 | 2.86 | 151.3 |
| 23 | B:SIA_1:O9(HO9) | A:ASN_248:OD1 | 3.9 | 2.80 | 153.8 |
| 24 | A:LYS_84:NZ(HZ2) | B:SIA_1:O1A | 3.8 | 2.82 | 155.3 |
| 25 | A:LYS_84:NZ(HZ1) | B:SIA_1:O1B | 3.8 | 2.82 | 155.3 |
| 26 | A:LYS_84:NZ(HZ3) | B:SIA_1:O1A | 3.8 | 2.82 | 155.6 |
| 27 | A:LYS_84:NZ(HZ2) | B:SIA_1:O1B | 3.6 | 2.83 | 155.2 |
| 28 | A:LYS_84:NZ(HZ3) | B:SIA_1:O1B | 3.6 | 2.83 | 155.3 |

**Extended Data Table 5. HKU1 N1 S1$^{A}$-ligand hydrogen bond analysis.**

H-bonds were identified based on the criteria given in Extended Data Fig. 23. Distances were calculated between heavy atoms, angles were measured between donor, H, and acceptor.

| Index | Donor | Acceptor | Population | Distance | Angle |
|---|---|---|---|---|---|
| 1 | B:SIA_2:N5(HN5) | A:LYS_80:O | 95.9 | 2.93 | 155.0 |
| 2 | A:THR_82:N(H) | B:SIA_2:O8 | 83.4 | 2.92 | 158.3 |
| 3 | A:LYS_80:NZ(HZ3) | B:SIA_2:O1B | 62.8 | 2.86 | 152.6 |
| 4 | A:THR_82:OG1(HG1) | B:SIA_2:O1B | 62.0 | 2.92 | 149.5 |
| 5 | A:LYS_80:NZ(HZ3) | B:SIA_2:O1A | 54.1 | 2.98 | 140.8 |
| 6 | B:SIA_2:O8(HO8) | A:TYR_84:O | 47.5 | 2.90 | 147.2 |
| 7 | A:THR_30:OG1(HG1) | B:SIA_2:O10 | 34.3 | 2.76 | 161.4 |
| 8 | A:THR_82:OG1(HG1) | B:SIA_2:O1A | 33.8 | 2.71 | 164.1 |
| 9 | A:ASN_26:ND2(HD22) | B:9AC_3:OA9 | 20.2 | 2.92 | 157.3 |
| 10 | A:SER_246:OG(HG) | B:SIA_2:O1A | 15.7 | 2.69 | 162.7 |
| 11 | B:SIA_1:N5(HN5) | A:THR_82:OG1 | 14.9 | 3.00 | 156.3 |
| 12 | B:SIA_1:O4(HO4) | A:ASN_248:O | 14.5 | 2.81 | 156.9 |
| 13 | A:ASN_243:ND2(HD22) | B:SIA_1:O1A | 9.6 | 2.88 | 161.5 |
| 14 | A:LYS_80:NZ(HZ2) | B:SIA_2:O1B | 8.6 | 2.84 | 151.3 |
| 15 | A:ASN_243:ND2(HD22) | B:SIA_1:O1B | 5.8 | 2.90 | 159.6 |
| 16 | A:ASN_26:ND2(HD21) | B:9AC_3:OA9 | 5.1 | 2.98 | 157.1 |
| 17 | A:THR_82:N(H) | B:SIA_2:O1B | 5.0 | 2.93 | 142.9 |
| 18 | B:SIA_2:O8(HO8) | A:THR_82:O | 4.9 | 2.86 | 144.0 |
| 19 | A:THR_31:OG1(HG1) | B:SIA_2:O10 | 4.8 | 2.80 | 155.2 |
| 20 | A:LYS_80:NZ(HZ1) | B:SIA_2:O1B | 4.4 | 2.85 | 151.0 |
| 21 | A:ASN_248:ND2(HD21) | B:SIA_1:O5N | 3.4 | 2.96 | 148.6 |
| 22 | B:SIA_1:O9(HO9) | A:4YB_3:O2N | 3.3 | 2.75 | 160.0 |
| 23 | A:ASN_248:ND2(HD22) | B:SIA_1:O1B | 3.2 | 2.87 | 158.2 |
| 24 | A:ASN_248:ND2(HD22) | B:SIA_1:O1A | 3.0 | 2.87 | 158.4 |

**Extended Data Table 6. Atom-atom contact count between HKU1 Caen1/N1 S1$^{A}$ and the disialoside ligand.**

The analysis was based on simulations starting from the *holo* state of Caen1 and N1 S1$^{A}$ domains in complex with the disialoside ligand with accumulated simulation times of 3 µs and 7 µs, respectively. Potential H-bonds between matching atom pairs were identified based on a heavy atom distance threshold of 3.2 Å. For hydrophobic contacts, a distance threshold of 4 Å was used for atoms not capable of hydrogen bonding. Salt bridges refer to distances smaller than 6 Å between Lys NZ/Arg CZ and C1 atoms of the ligand. Favourable contacts (the sum of H-bond, hydrophobic and salt bridge contacts) per ligand residue are also shown in Extended Data Fig. 27.

| | **HKU1-A Caen1** | | **HKU1-A N1** | |
|---|---|---|---|---|
| **ligand-receptor contacts** | **mean** | **std** | **mean** | **std** |
| total number of contacts | 76.0 | 12.9 | 73.8 | 15.5 |
| total H-bond contacts | 6.9 | 1.7 | 8.1 | 1.9 |
| total hydrophobic contacts | 17.4 | 3.9 | 14.5 | 4.4 |
| total salt bridges | 1.4 | 0.5 | 1.0 | 0.2 |
| total favourable contacts SIA2 | 14.0 | 3.1 | 14.7 | 3.6 |
| H-bond contacts SIA2 | 5.3 | 1.3 | 6.6 | 1.4 |
| hydrophobic contacts SIA2 | 7.7 | 2.8 | 7.1 | 3.1 |
| salt bridges SIA2 | 1.0 | 0.2 | 1.0 | 0.2 |
| total 9-O-Ac favourable contacts | 7.5 | 2.4 | 5.3 | 2.2 |
| H-bond contacts 9-O-Ac | 0.6 | 0.5 | 0.3 | 0.5 |
| hydrophobic contacts 9-O-Ac | 6.9 | 2.2 | 5.0 | 2.1 |
| total SIA1 favourable contacts | 4.2 | 2.4 | 3.3 | 2.6 |
| H-bond contacts SIA1 | 1.0 | 1.0 | 1.1 | 1.1 |
| hydrophobic contacts SIA1 | 2.7 | 1.9 | 2.1 | 2.0 |
| salt bridges SIA1 | 0.5 | 0.5 | 0.0 | 0.1 |

**Supplementary Video 1.**

Morph between the *apo*, *apo* with ligand placed into the binding site, closed *holo*, *holo* 1-up and *holo* 3-up atomic models.

**Supplementary Video 2.**

$S1^{A}$ domain from one protomer (light grey) and the counterclockwise neighbouring $S1^{B}$ domain (orange), morphing from *apo* to closed *holo* to the *holo* up state. Note that the inward subdomain rotation in $S1^{A}$ changes the $S1^{A}$-$S1^{B}$ interface substantially.

**Supplementary Video 3.**

Morph between the *apo* and closed *holo* locally refined maps. The maps were aligned on the $S1^{A1}$ sub-domain.

**Supplementary Video 4.**

Close-up of the disialoside binding pocket in $S1^{A}$, morphing from *apo* to closed *holo*, coloured as in Extended Data Fig. 4a/b and aligned on the shown binding site.

**Supplementary Video 5.**

Visualisation of the inward wedging subdomain rotation of $S1^{A1}$ with respect to $S1^{A2}$ upon disialoside ligand binding (again morphing from *apo* to closed *holo*), coloured as in Extended Data Fig. 4c/d.

**Supplementary Video 6.**

Visualisation of the high-resolution MD-derived pseudo-density model of the disialoside in the HKU1 binding pocket.

**Supplementary Video 7.**

Conformational change in e1 as predicted by MD simulations after docking of the disialoside ligand into the cryo-EM *apo* model. The transition shown corresponds to the one depicted in Fig. 5b and was obtained from a 1 us MD simulation.
